# TridentTCR: Predicting T Cell Receptor Specificity for Antigens and Autoimmune-Related Targets via a Topology-Aware Graph and Large Language Model

**DOI:** 10.1101/2025.06.27.661632

**Authors:** Rui Niu, Xiaoying Kong, Jiren Zhou, Yanli Li, Xuequn Shang

## Abstract

T cells recognize and eliminate diseased cells by binding their T cell receptors (TCRs) to short endogenous peptides presented on the cell surface, commonly referred to as antigens. Accurate prediction of antigen–TCR specificity is essential for advancing T cell-based immunotherapies while minimizing autoimmune toxicity. However, current computational methods struggle to generalize across diverse TCR repertoires and reliably differentiate therapeutic antigens from autoimmune-related antigens (arAgs). Here, we introduce TridentTCR, a computational framework integrating a pretrained large language model (LLM) for high-fidelity sequence embedding with a topology-aware graph neural network to capture structural insights from the antigen–TCR interaction network. TridentTCR significantly advances TCR specificity prediction by enabling robust trinary classification, distinguishing TCR interactions involving general antigens, arAgs, and non-binding. Additionally, TridentTCR outperforms existing state-of-the-art models in binary classification (binding vs. non-binding). Validation on independent clinical datasets demonstrated strong generalization capability for unseen TCRs and antigens. In a disease-specific context, TridentTCR identifies notable cross-reactivity between ZnT8_186-194_-specific TCRs and a mimotope from *Bacteroides stercoris*, suggesting that T cell cross-reactivity may contribute to the initiation of type 1 diabetes. Furthermore, we introduced a quantitative metric, antigenic immune response entropy (AIRE), which leverages TridentTCR predictions alongside clonotype frequencies and repertoire diversity to precisely quantify antigen-specific immune responses from single-cell profiling data. Collectively, TridentTCR provides an interpretable and clinically relevant tool, enabling improved understanding of TCR specificity, cross-reactivity, and off-target autoimmune risks in clinical immunotherapy.

## Introduction

T cells initiate immune responses to effectively eliminate viruses and tumours through antigen-specific recognition. Specifically, they use T cell receptors (TCRs) to recognize short endogenous peptides—commonly referred to as antigens—presented by major histocompatibility complex (MHC) molecules on the surface of diseased cells, thereby eliciting antigen-specific T cell responses. Leveraging the specificity of antigen-TCR binding, targeted tumour immunotherapies have significantly reshaped cancer treatment over the past decade. However, multiple studies have shown that these therapies can induce toxicities due to autoimmune responses triggered by T cell cross-reactivity, where a single TCR is capable of recognizing multiple antigens^1,2,3,4^. A precise and systematic characterization of TCR recognition of targeting antigens and autoimmune-related antigens (arAgs) will advance our understanding of T cell specificity and support the development of safer and more effective immunotherapies.

One of the primary methods for identifying TCRs that bind a specific antigen is to design a peptide-major histocompatibility complex class I (pMHC-I) tetramer for a known peptide sequence, use it to enrich T cells whose TCRs recognize that complex, and then sequence those TCRs. Although tetramer-based assays offer high specificity, their application is limited by the substantial costs of generating tetramers. Research advances in this field, coupled with the evolution of sequencing technologies, have generated extensive immune repertoire data now publicly available in various databases. These developments create new opportunities for computational methods to predict antigen-TCR interactions.

With the availability of large-scale immune repertoire data, several computational methods have been developed for predicting antigen–TCR interactions ^5-7^. Early unsupervised methods such as TCRdist, iSMART, and GIANA clustered TCR sequences based on similarity to infer antigen-specific recognition patterns ^8-10^. With the growing accumulation of high-quality antigen–TCR interaction data, supervised computational methods have rapidly advanced. These supervised approaches can be broadly classified as biologically informed encoders and pretrained encoders based on their feature-encoding strategies. Biologically informed encoders include NetTCR-2.0, which employs the BLOSUM50 substitution matrix, and PanPep, which utilizes Atchley factors to encode antigen and TCR sequences ^5,11^. Pretrained encoders, such as those in ERGO-II, pMTnet, and HeteroTCR, employ neural network architectures including long short-term memory (LSTM), autoencoders, and convolutional neural networks (CNNs) ^6,12,13^. These models are pretrained on extensive antigen and TCR sequence corpora to derive high-dimensional embeddings, which are further refined by transfer learning to enhance their representational power. Although these machine learning approaches have substantially improved feature extraction and boosted prediction accuracy, the clinical efficiency of computational algorithms lies at whether they could overcome the significant diversity of TCR repertoires among individuals? High-throughput sequencing of T cells from the peripheral blood of healthy adults finds ∼ 5 to 6 million TCRs per individual, but only a very small fraction of these is shared between donors— with shared TCRs account for < 1% of peripheral blood CD8+ T cells even between siblings ^14^. While both feature-encoding strategies have achieved a certain level of success, these relatively shallow encoding methods are confined by the narrow diversity of their training data and thus learn primarily common amino-acid motifs and conserved residues. As a result, when faced with novel sequence fragments or uncommon combinations from individual TCR repertoires, the models cannot reliably project them into the learned embedding space.

Furthermore, T cells in the immune system exhibit cross-reactivity, where a single TCR can recognize multiple antigens^1^. Conversely, studies have also shown that T cells often utilize multiple distinct TCRs to recognize the same antigen^15^. This many-to-many relationship naturally maps onto a heterogeneous bipartite graph, with antigens and TCRs as node types and their binding events as edges. Exploiting this structure with graph, HeteroTCR has demonstrated superior performance in antigen–TCR interaction prediction^8^. Such models offer a powerful route for selecting T-cell populations most likely to target tumour antigens, yet their clinical utility is limited by residual uncertainties in specificity arising from TCR cross-reactivity. Immune-checkpoint inhibitors (ICIs) reactivate anti-tumour T cells by blocking inhibitory receptor–ligand pairs and are now a mainstay of cancer immunotherapy^16^. Nevertheless, cross-reactive TCRs can also bind self-derived peptides, leading to immune-related adverse events such as vitiligo in melanoma patients treated with ICIs^2,3^. Accurately delineating targeting interactions from those involving arAg therefore remains an urgent objective for antigen-TCR interaction prediction models.

In this work, we integrate complementary sequence and topological information to enhance predictions of TCR recognition specificity, particularly addressing the critical challenge of distinguishing TCR interactions with general antigens from arAgs, broadly defined as antigens capable of aberrantly activating immune responses. To address current limitations in generalizing across diverse TCR repertoires and unseen antigens, we introduce TridentTCR, a robust computational framework that synergistically integrates a pretrained large language model (LLM) with a topology-aware graph neural network (GNN). Specifically, TridentTCR employs the pretrained LLM to encode amino-acid sequences of antigens and TCRs into high-dimensional embeddings, which are subsequently integrated within a geometric multi-view aggregation scheme that captures both sequence-level specificity and network-level relational context. Comprehensive evaluation demonstrates TridentTCR’s superior performance in a novel trinary classification setting (general antigen–TCR binding vs. arAg–TCR binding vs. non-binding). Additionally, a binary classification variant, distinguishing solely between binding and non-binding interactions, outperforms existing state-of-the-art approaches. Crucially, when evaluated on clinical datasets containing numerous previously unseen TCRs and antigens, TridentTCR maintains robust performance, underscoring its ability to overcome the inherent diversity of individual TCR repertoires. To further quantify patient-specific immune responses, we propose a quantitative metric termed antigenic immune response entropy (AIRE). AIRE integrates TridentTCR-predicted recognition probabilities, T cell clonotype frequencies, and TCR repertoire diversity to precisely characterize antigen-specific immune responses at the individual level. Applied to human single-cell immune-sequencing data, AIRE effectively distinguishes disease-associated immune states across multiple donors and pathogens. To our knowledge, TridentTCR represents the first computational approach explicitly designed to differentiate TCR specificity toward general antigens versus autoimmune-related antigens, providing critical new insights into TCR cross-reactivity and off-target risks in clinical T cell-based immunotherapies.

## Results

### The framework of TridentTCR

TridentTCR is a robust framework designed to distinguish TCR specificity toward general antigens and arAgs by integrating sequence-based embeddings with a geometry-aware graph learning approach. Conceptually, TridentTCR comprises three core stages (Fig. 1). First, numeric embeddings of antigen and TCR sequences are extracted using ESM-2^17^. Latent neighborhoods are then constructed by identifying the similarities and geometric relationships within the antigen or TCR embedding spaces. Second, we build a heterogeneous graph that integrates nodes for antigen and TCR embeddings and edges reflecting both cross-type interactions and latent intra-type similarities. The multi-view geometric graph learning framework is employed to propagate information across both interaction and latent similarity views. Finally, we implement a downstream multi-class multi-layer perceptron (MLP) classifier to distinguish among three interaction modes: antigen–TCR binding, arAg–TCR binding, and no interaction.

**Fig. 1.**
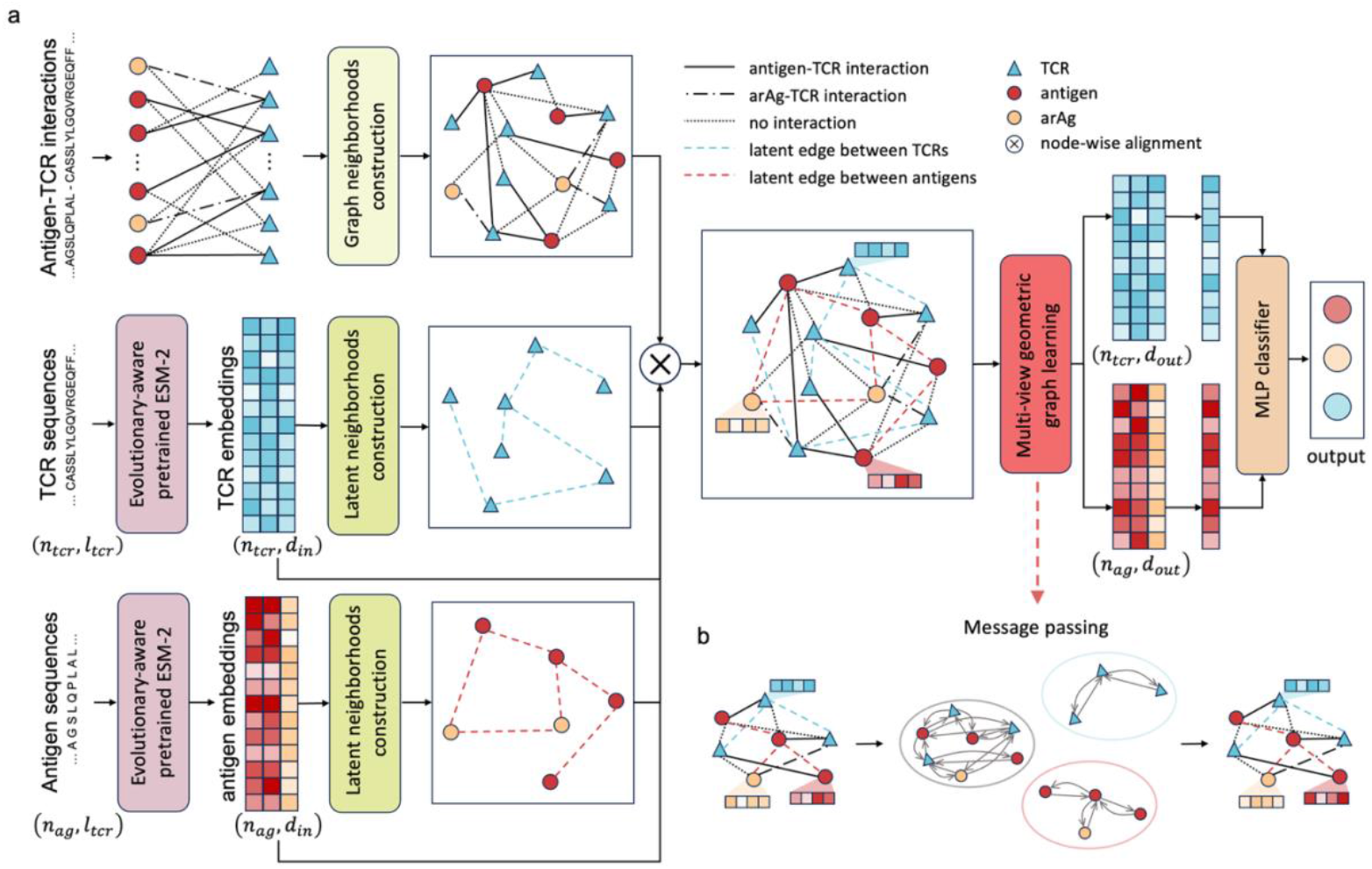
Overview of the TridentTCR framework for predicting TCR binding specificity to antigens and arAgs. **a**, Structure of the TridentTCR model. The observed interaction network defines three types of TCR binding modes: antigen–TCR (solid), arAg–TCR (dash-dotted), and no interaction (dotted), which are used to construct graph neighborhoods. Input antigen and TCR sequences are encoded using the pre-trained ESM-2 model to generate high-dimensional embeddings. Latent neighborhoods are independently constructed for TCRs (blue dashed lines) and antigens (red dashed lines) based on geometric proximity in the embedding space. Graph and latent edges are integrated via node-wise alignment, forming a heterogeneous graph where nodes represent TCRs or antigens and edges denote their observed or geometric relationships. Node features are updated through a multi-view geometric graph learning framework, and an MLP classifier is subsequently used to predict the probability of each binding mode. **b**, Message passing in TridentTCR. The multi-view geometric graph learning framework propagates information across both interaction and latent views, allowing the model to jointly capture interaction topology and sequence similarity to refine node representations.

To encode the amino acid sequences of TCRs and antigens, we employed ESM-2, a transformer-based LLM pretrained on massive protein corpora and comprising more than 650 million parameters. For TCR sequences, we utilized the complementarity determining region 3 (CDR3) region of the TCR β chain (TCRβ), given its critical role in antigen specificity determination^18^. Each antigen and TCR sequence is thus represented as a 1280-dimensional embedding, capturing rich contextual and evolutionary representations. Subsequently, these embeddings are projected into latent spaces separately for antigens and TCRs, in which intra-type of similarity edges (antigen– antigen and TCR–TCR) are dynamically defined based on geometric proximity. For each pair of nodes connected by latent edges, their relative geometric relationships are computed for subsequent message aggregation.

We employed a biologically informed heterogeneous graph to represent the interaction network includes TCRs, antigens, and arAgs. In this graph, two types of nodes represent antigen and TCR embeddings, while edges are categorized into two complementary views. The interaction view consists of cross-type edges, capturing experimentally validated binding relationships between TCRs and their cognate antigens or arAgs. The latent view consists of intra-type similarity edges, reflecting antigen–antigen similarities based on shared recognition by TCRs, and TCR–TCR similarities arising from common antigen recognition patterns. Thus, the graph structure simultaneously encodes molecular specificity derived from direct binding interactions and broader structural contexts provided by similarity-driven latent relationships. To aggregate information across these complementary views, we adopted a multi-view geometric graph learning framework. This dual-view aggregation enables the model to simultaneously capture biologically meaningful interactions between distinct node types and intrinsic geometric similarities within the same node type, thereby effectively updating node representations (Fig. 1b).

Finally, leveraging the refined embeddings from graph learning, we employed a MLP classifier. The concatenated antigen–TCR embeddings are passed through three linear layers, producing output probabilities for three interaction states: antigen–TCR binding, arAg–TCR binding, and non-binding interactions. Each output neuron yields a continuous probability between 0 and 1, allowing for robust trinary classification of TCR specificity.

### TridentTCR accurately predicts antigen-TCR and arAg-TCR specificities

We collected antigen–TCR interaction data from public databases including IEDB, VDJdb, and McPAS-TCR^19-21^. Where available, database-provided quality metrics were used to filter records, retaining only high-confidence interaction pairs. After preprocessing, we obtained a total of 41,732 high-quality binding pairs; detailed preprocessing steps are provided in the Methods. Additionally, antigen annotations from IEDB and McPAS-TCR were employed to identify antigens associated with autoimmune diseases, which we labeled as arAgs. Within our benchmark dataset, interactions involving arAgs were labeled as arAg–TCR pairs (N = 11,054), while the remaining interactions were classified as antigen–TCR pairs (N = 30,678). We trained our model using five-fold cross-validation, randomly partitioning data into five equally sized subsets. In each iteration, negative samples equal in number to positive pairs were generated by randomly mismatching TCR and antigen sequences within each fold. The model performance was evaluated iteratively on each subset in a rotating manner.

In T cell-based immunotherapies, false-positive predictions in antigen–TCR recognition—cases where a TCR is incorrectly predicted to bind a particular antigen—often pose greater risks than false negatives, which merely overlook potential therapeutic TCRs. For instance, a TCR mistakenly identified as capable of recognizing a tumor antigen may be used to engineer therapeutic T cells (such as T cells in TCR-T or CAR-T therapies). If this TCR does not genuinely bind the intended antigen or instead cross-reacts with similar peptides presented on healthy cells, severe on-target, off-tumour toxicity can occur^22^. This phenomenon leads to unintended damage to healthy tissues, potentially causing irreversible injury or life-threatening autoimmune responses. To mitigate this risk during training, we implemented a hybrid loss function specifically designed to impose stronger penalties on false-positive predictions. This loss function reduces gradient contributions from easily classified samples and amplifies the loss for challenging samples, thereby focusing model attention on cases that are difficult to distinguish, near decision boundaries, or underrepresented (Methods).

To evaluate the overall performance of TridentTCR on the three-class prediction task, we utilized macro area under the receiver operating characteristic curve (AUROC), macro area under the precision-recall curve (AUPRC), and macro accuracy as performance metrics. As shown in Fig. 2a, our five-fold cross-validation experiments yielded an AUROC of 0.904 ± 0.006, an AUPRC of 0.854 ± 0.010, and an accuracy of 0.743 ± 0.009. Furthermore, we plotted one-vs-rest AUROC curves for each class individually, achieving scores of 0.893 ± 0.006 for antigen–TCR interactions and 0.979 ± 0.003 for arAg–TCR interactions (Fig. 2b). To verify that TridentTCR genuinely learned predictive binding features, we further evaluated the stability of model performance under varying thresholds of TCR similarity (Fig. 2c). We partitioned the benchmark dataset into 80% training data and 20% test data. For each TCR in the test set, we calculated its euclidean distance in feature space to all TCRs in the training set, retaining only the minimum distance (see Supplementary Information). These minimum distances ranged from 0.0005 to 0.854, exhibiting a clear long-tailed distribution (Supplementary Fig. 1). We iteratively removed the top 10% most similar pairs based on the minimum distance, and re-evaluated the performance using AUROC, AUPRC, and accuracy. Across all iterations, even after removing up to 80% or 90% of the most similar pairs, all metrics remained consistent. This indicates that TridentTCR achieves robust generalization, rather than relying on highly similar or near-duplicate sequences to maintain predictive accuracy. Collectively, these results demonstrate that TridentTCR robustly distinguishes between TCRs targeting general antigens, arAgs, or neither in a trinary classification setting.

**Fig. 2.**
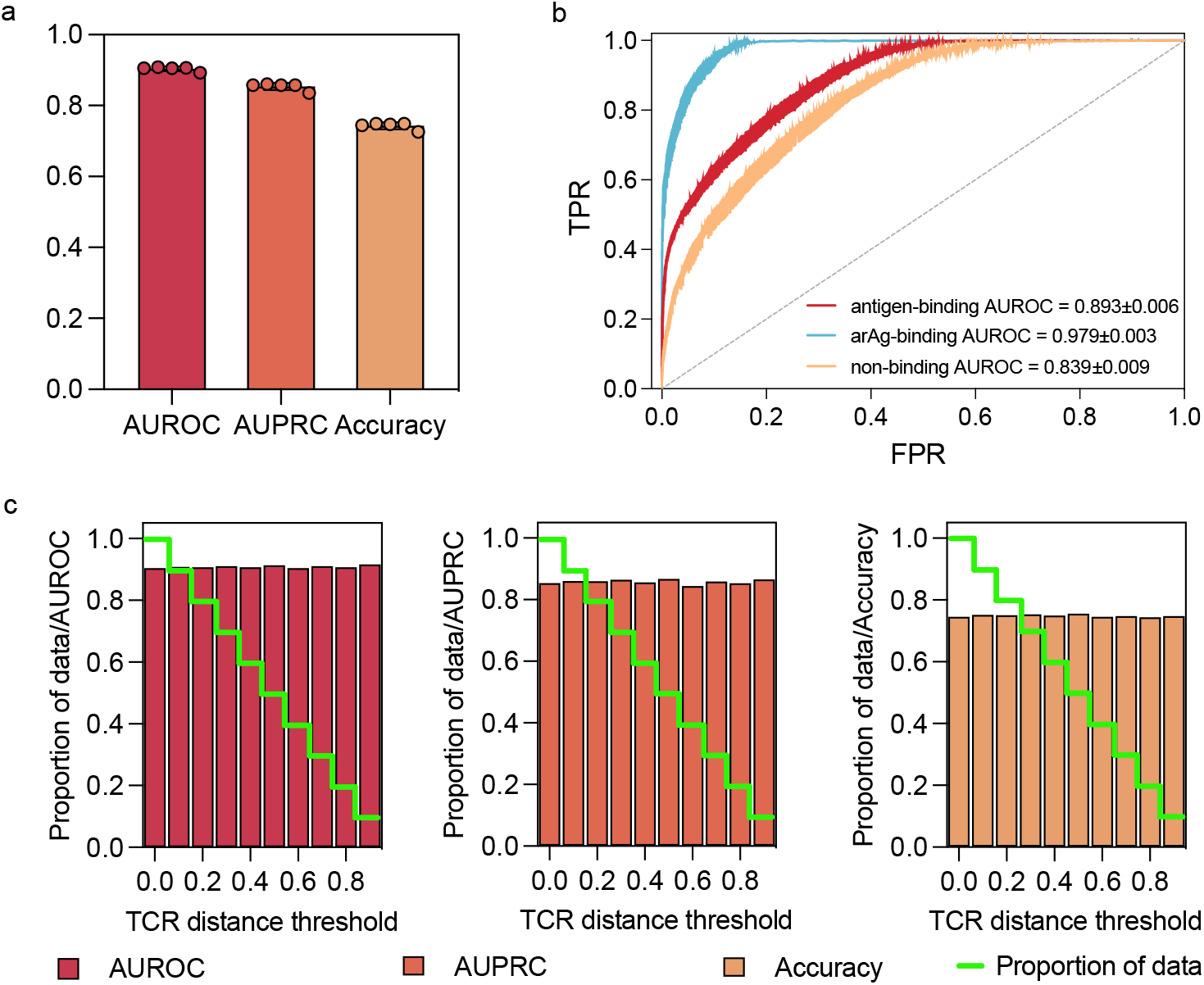
TridentTCR accurately predicts antigen and arAg binding specificity across diverse TCRs. **a**, Macro AUROC, AUPRC, and accuracy of TridentTCR evaluated on the benchmark dataset using five-fold cross-validation. **b**, One-vs-rest AUROC curves for antigen-binding, arAg-binding, and non-binding predictions. Solid lines represent the mean AUROC across five cross-validation folds, and shaded regions indicate standard deviations. **c**, Robustness analysis of TridentTCR under increasing thresholds of TCR dissimilarity. Test TCRs were ranked by their minimum euclidean distance to the training TCRs, and the most similar 10% of pairs were progressively removed. Performance metrics are plotted for the remaining data subsets, while the green line shows the proportion of data retained at each step.

**Fig. 3.**
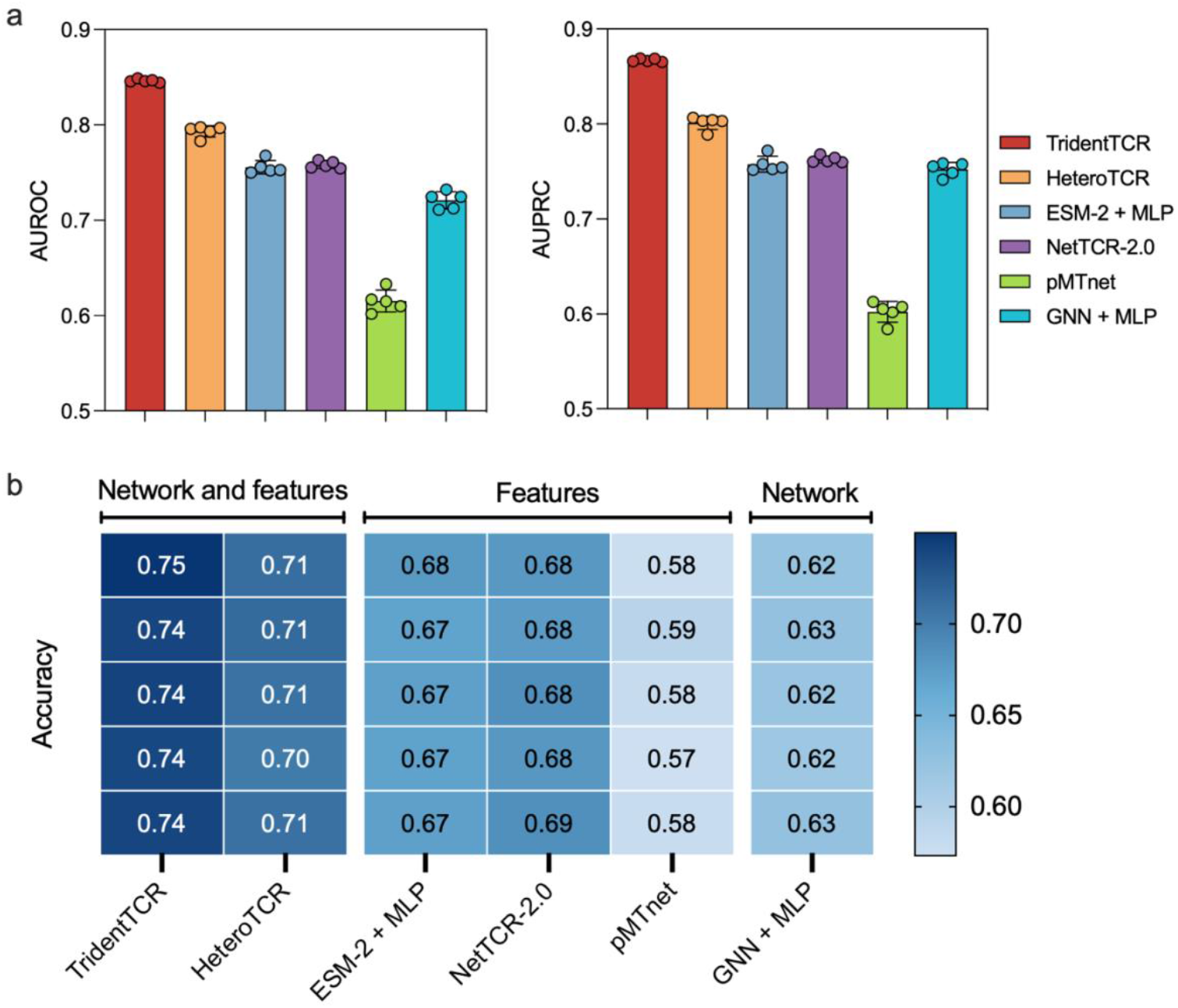
Comparison of TridentTCR with baseline models in predicting TCR–antigen interactions. **a**, AUROC and AUPRC of TridentTCR and five baseline models on a benchmark dataset using five-fold cross-validation. Antigen–TCR and arAg–TCR interaction pairs were merged and treated as positive samples. **b**, Heatmap of prediction accuracy for all methods across five folds, grouped by the type of input information used: combined sequence and network data, sequence features only, or network topology based.

### Performance comparison with other methods

As TridentTCR is currently the only model distinguishing TCR specificity between antigens and arAgs, direct comparisons with existing methods—which exclusively predict antigen–TCR binding—are not feasible. To facilitate such comparisons, we introduced a binary-classification variant of TridentTCR, modified specifically to predict antigen–TCR interactions without differentiating antigens from arAgs (Supplementary Information). Accordingly, antigen–TCR and arAg–TCR interaction pairs were combined into a unified positive dataset (N = 41,732). For all models evaluated, we employed five-fold cross-validation, generating an equal number of negative samples by randomly mismatching positive pairs within each fold.

To demonstrate the advantages of integrating diverse data types, we benchmarked TridentTCR against methods relying on only single data modalities—either exclusively sequence-based features or network topology alone. We selected two mainstream feature-only models for comparison: NetTCR-2.0 and pMTnet, both adopting pair-based strategies wherein each antigen– TCR pair is treated as an independent sample^5,6^. NetTCR-2.0 encodes antigen and TCR sequences using the biologically informed BLOSUM50 matrix, subsequently training a shallow CNN for prediction. In contrast, pMTnet employs a transfer-learning strategy, encoding TCR sequences via an autoencoder pretrained on 243,747 TCRβ CDR3 sequences, and antigen sequences via a LSTM model pretrained on 172,422 pMHC-I sequences, before ultimately training a fully connected deep neural network for TCR recognition prediction. Notably, for NetTCR-2.0, we exclusively employed the TCRβ CDR3 region as the TCR input, given its pivotal role in antigen recognition. Although pMTnet originally incorporates additional MHC-I molecule information, we focused exclusively on antigen–TCR interactions in this study; thus, we adapted pMTnet by substituting antigen sequences for the original pMHC-I inputs, while still leveraging its pretrained encoders. We further designed a feature-based model to encode immune sequences by ESM-2 then classified by an MLP classifier, to directly assess differences between our feature-only variant (TridentTCR without topological information) and established methods such as NetTCR-2.0 and pMTnet. In parallel, we also implemented a network-based approach integrating the GNN module with an MLP classifier. This method employed the basic BLOSUM50 encoding as input rather than ESM-2 embeddings, relied solely on GNN-derived latent network features, and concatenated antigen and TCR features before classification with the MLP. Additionally, we benchmarked TridentTCR against HeteroTCR, a recently developed method integrating both sequence and network features^13^. HeteroTCR encodes immune sequences using a pretrained CNN and employs a three-layer GNN to learn network topology. Conceptually, HeteroTCR is most similar to TridentTCR, as it incorporates precisely the same input feature types, yet it differs fundamentally in its underlying machine learning architecture.

As shown in Fig. 3a, the binary-classification variant of TridentTCR achieves an AUROC of 0.846 ± 0.002 and an AUPRC of 0.867 ± 0.002 in predicting antigen–TCR interactions. When grouping the models based on the type of data used (Fig. 3b), we observed that models integrating both sequence features and network topology outperformed those relying exclusively on sequence features, which, in turn, surpassed models solely leveraging network topology (i.e., the combination of a GCN module with an MLP classifier). Among feature-only methods, the model combining ESM-2 embeddings and an MLP classifier significantly outperformed pMTnet by an average accuracy margin of 9.2%, directly validating the efficacy of ESM-2 as a feature encoder. Overall, TridentTCR consistently outperformed all competing models in the binary classification setting, exceeding the second-best approach by 5.3% in AUROC, 6.6% in AUPRC, and 3.3% in accuracy, on average.

### TridentTCR benefits from different data representations

To determine which input modality contributes most to TridentTCR’s predictive power, we began by comparing the pretrained ESM-2 model with other encoding schemes for representing antigen-specific TCRs. We compiled a non-redundant set of TCR sequences from VDJdb, together with their cognate antigens and epitope-species annotations. The resulting dataset comprised 8,506 unique TCRβ CDR3 sequences spanning 19 epitope species. Each sequence was encoded using either BLOSUM50 or ESM-2. The resulting embeddings were projected using UMAP (Fig. 4a), with each point coloured by epitope species. BLOSUM50 produced diffuse, overlapping clusters with no clear species-level separation, implying that the substitution matrix captures broad biochemical similarity but cannot resolve the fine-grained sequence difference that underlie TCR specificity. By contrast, ESM-2 formed tight, well-segregated clusters and even resolved subtle differences among TCRs targeting the same epitope species. Further comparisons with HeteroTCR’s CNN-based encoder and pMTnet’s autoencoder (Supplementary Fig. 2) confirmed ESM-2’s superior latent structure. Quantitative evaluation using Davies–Bouldin index and intra-class metrics (Supplementary Tables 2–4) showed ESM-2 consistently outperformed other methods, highlighting its capacity to capture immunologically relevant sequence features that conventional encodings miss.

**Fig. 4.**
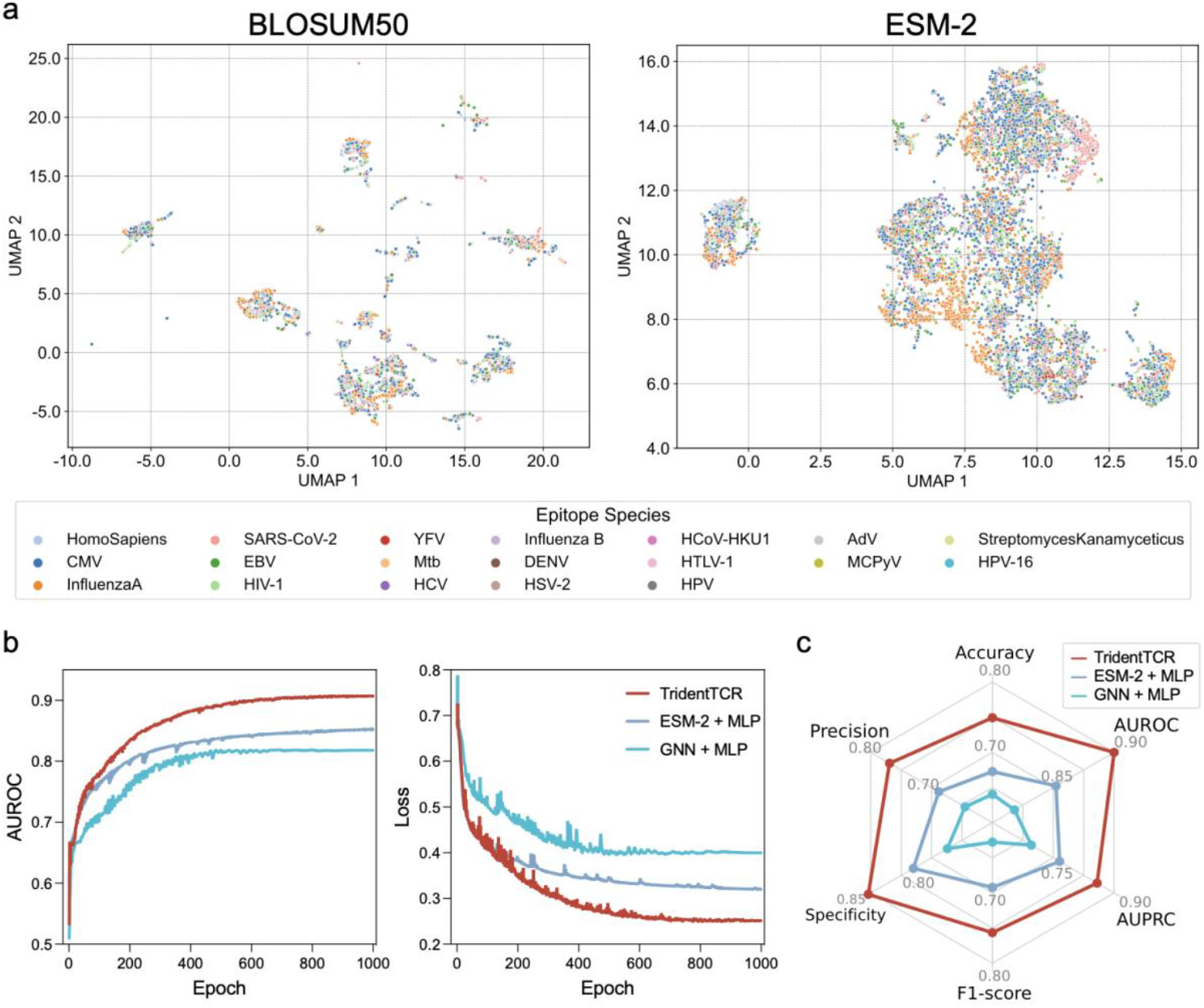
Integration of informative sequence features and network topology enhances the predictive performance of TridentTCR. **a**, Comparison of representations of TCRs from VDJdb, learned using either a biologically informed encoder (BLOSUM50) or a pre-trained large language model (ESM-2). UMAP projections of the resulting embeddings are shown, with each point colored by its recognized epitope species. **b**, Training dynamics of the full TridentTCR model compared with two ablated variants: a feature-only model (ESM-2 + MLP) and a network-based model (GNN + MLP, using BLOSUM50 for minimal sequence features). Left: macro AUROC over training epochs. Right: loss curves over the same training period. **c**, Comparative evaluation of the three models across six performance metrics: macro accuracy, AUROC, AUPRC, F1-score, specificity, and precision. The radar plot highlights TridentTCR’s superior performance across all dimensions. Perturbing either the sequence features or the network structure resulted in significant performance declines across all six metrics (paired t-test, *p* < 0.001); detailed comparisons are shown in Supplementary Fig. 3.

We subsequently performed ablation experiments on the three-class dataset, and compared the modified model with the original TridentTCR. To assess the contribution of network topology information, we designed a feature-only variant that combines ESM-2 embeddings with an MLP classifier. In parallel, we implemented a network-based variant that uses BLOSUM50 in place of ESM-2 to provide minimal sequence feature information, as described previously. The output layer of both models was modified to produce three continuous values, each ranging from 0 to 1, representing the predicted probabilities of antigen–TCR binding, arAg–TCR binding, or no interaction. All three models were trained on 80% of the data and tested on the remaining 20%, with performance evaluated using six metrics including macro AUROC and macro AUPRC (Fig. 4b–c). The results indicate that both components contribute substantially: removing the GNN or the LLM module led to a 5.5% and 8.9% drop in AUROC, respectively (Fig. 4b). As shown in Fig. 4c, perturbing either the sequence features or the network structure resulted in significant performance declines across all six metrics compared to TridentTCR (paired t-test, *p* < 0.001), with details provided in Supplementary Fig. 3. Notably, the LLM contributed more—replacing ESM-2 with BLOSUM50 caused a greater performance drop than removing the network, confirming that high-fidelity sequence representations are indispensable for accurate TCR specificity prediction. Collectively, these results demonstrate that TridentTCR benefits from the integration of sequence embeddings and topological information, with high-fidelity sequence representations contributing more significantly to predictive performance.

### TridentTCR enables cross-individual generalization

The remarkable diversity of TCR repertoires across individuals has long posed a major challenge to the accurate prediction of TCR binding specificity. This highlights the importance of evaluating whether computational models can generalize across individuals despite such repertoire variability. To systematically assess the generalizability of TridentTCR, we compiled three independent external test sets from previously published studies, thereby also demonstrating the model’s applicability in clinically relevant contexts. The first test dataset, derived from 10x Genomics, integrates single-cell immune profiling with tetramer staining to evaluate the specificity of approximately 150,000 CD8^+^ T cells against 44 distinct antigen-specific tetramers across four donors^23^. The second dataset includes 44 distinct TCRs from a single donor, specific to four viral antigens: Influenza A M1_58-66_ (GILGFVFTL), Influenza A PA_46-54_ (FMYSDFHFI), EBV BMLF1_280-288_ (GLCTLVAML), and HCMV pp65_495-503_ (NLVPMVATV) ^6^. The third dataset focuses on autoimmunity, comprising 14 autoreactive TCRs from 12 donors, all recognizing a type 1 diabetes (T1D)– associated antigen ZnT8_186-194_ (VAANIVLTV)^24^.

#### TridentTCR predict the antigen-specific T cell clonotypes for unseen TCRs

We first analyzed the datasets from 10x Genomics. For each donor, we focused on the top 1% of T cell clonotypes ranked by clonal frequency. Specifically, for each clonotype, we quantified the unique molecular identifier (UMI) counts associated with all antigen-specific tetramers, and retained only those clonotypes with a UMI count greater than 10 for at least one tetramer. We refer to the set of TCRs corresponding to these clonotypes as the antigen-specific TCR repertoire (details provided in Supplementary Information). To assess overlap in antigen-specific TCR repertoires, we first quantified clonotype sharing based on TCRβ CDR3 sequence. As shown in Fig. 5a (left), Donor1 and Donor2 shared a few clonotypes, while Donor3 and 4 showed little to no overlap with others. Overall, the extent of sequence-based repertoire overlap was limited, indicating substantial inter-individual diversity, consistent with previous findings^14^. We then examined repertoire overlap from a functional perspective by aggregating clonotypes according to their pMHC-I specificity (i.e., recognition of the same pMHC-I complex). As shown in Fig. 5a (right), the degree of overlap increased substantially under this criterion, with every donor pair sharing clonotypes specific to at least one common pMHC-I. To further explore this phenomenon, we encoded antigen-specific TCRs from all four donors by ESM-2 and projected the embeddings into a shared UMAP embedding (Fig. 5b). The resulting distribution was highly intermixed, lacking distinct donor-specific clusters. This observation supports the notion of shared antigen specificity despite sequence-level dissimilarity across individuals.

**Fig. 5.**
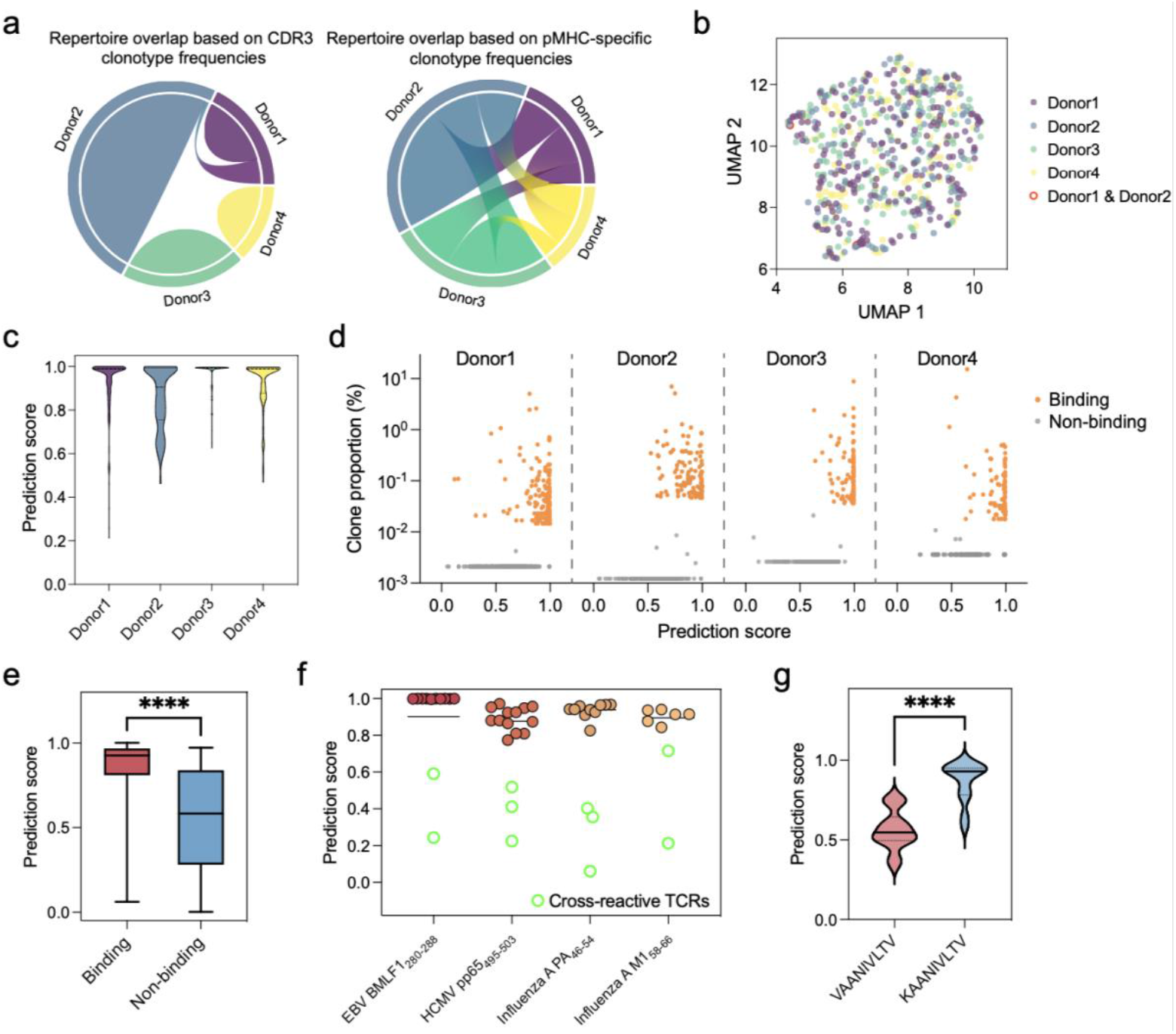
Validation of TridentTCR using independent external datasets. **a**, Antigen-specific repertoire overlap across four donors from 10x Genomics dataset, calculated based on CDR3β sequence similarity (left) and pMHC specificity (right). Chord colors indicate donor identity. **b**, UMAP projection of ESM-2 encoded antigen-specific TCRs from four donors. Each point is colored by donor, with shared TCRs between donor1 and donor2 highlighted as red open circles. **c**, Truncated violin plots showing the distribution of predicted binding scores for true TCR-antigen pairs from donor1 (n = 213), donor2 (n = 135), donor3 (n = 150), and donor4 (n = 106). Plots indicate medians (centres), interquartile ranges (hinges), and 1.5 times the interquartile ranges as whiskers. **d**, Predicted binding scores and clonal proportions for binding (orange) and non-binding (grey) pairs across four donors. High-scoring binding clonotypes tended to exhibit greater clonal expansion. **e**, Comparison of predicted binding scores between true binding pairs (red) and constructed non-binding pairs (blue) from a donor with prior viral infections and lung cancer. Binding pairs showed significantly higher scores (unpaired t-test, **** *p* < 0.0001). **f**, Predicted binding scores for 44 TCRs targeting four viral antigens. Each dot denotes a TCR– antigen pair. Green open circles highlight cross-reactive TCRs recognizing multiple antigens, which consistently showed lower predicted scores, indicating reduced binding specificity. **g**, Predicted binding score distributions of autoimmune-reactive TCRs recognizing T1D-associated antigen ZnT8_186-194_ (VAANIVLTV) and homologous microbial antigen from *B. stercoris* (KAANIVLTV). KAANIVLTV pairs exhibited significantly higher scores than VAANIVLTV (unpaired t-test, **** *p* < 0.0001).

For each T cell clonotype, we computed its minimum Euclidean distance to TCRs in the benchmark dataset. As shown in Supplementary Fig. 4a, over 60% of antigen-specific TCRs from each donor exhibited a minimum distance greater than 0.1 from any TCR in the benchmark dataset, indicating limited sequence similarity. To evaluate the performance of TridentTCR on these unseen TCRs, we used the five models trained via five-fold cross-validation on the benchmark dataset to independently predict the binding scores of antigen–TCR pairs and averaged the results to minimize the influence of data imbalance (Fig. 5c). To further assess whether TridentTCR can distinguish true binding from non-binding events, we constructed donor-specific negative samples. For each donor, we first excluded the top 1% of T cell clonotypes ranked by clonal frequency. Then, from the remaining repertoire, we removed all clonotypes with UMI counts >0 for any tetramer, and randomly sampled an equal number of clonotypes to match the antigen-specific TCRs in that donor. We also computed the minimum Euclidean distances from these negative TCRs to the benchmark TCRs. As shown in Supplementary Fig. 4a, the negative TCRs were overall more distant from the benchmark dataset than the antigen-specific TCRs. For each donor, we grouped the antigen–TCR binding pairs by antigen and sampled an equal number of negative TCRs for each antigen to construct non-binding antigen–TCR pairs. We then applied the five cross-validated models to predict the binding probabilities of both binding and non-binding pairs, and averaged the predictions to obtain the final binding scores. For each donor, we visualized the predicted binding scores alongside clonal proportions for all clonotypes (Fig. 5d). The prediction score distributions for true binding and non-binding pairs were clearly distinct. Antigen-specific clonotypes with higher prediction scores also tended to exhibit stronger clonal expansion, whereas non-reactive clonotypes showed both low prediction scores and limited expansion. An unpaired t-test further confirmed that the prediction scores of binding pairs were significantly higher than those of non-binding pairs (*p* < 0.0001), supporting the robustness of TridentTCR’s performance on unseen TCRs (Supplementary Fig. 5). However, T cell clonal expansion is highly context dependent. While clonotypes with higher predicted binding scores tended to expand more, the expansion levels across antigen-specific TCRs were not always predictable. This may be due to unmeasured differences in TCR–antigen affinity and inter-clonal competition^25^.

We next evaluated TridentTCR on a dataset collected from a donor with a history of Influenza, EBV, and HCMV infections, and currently diagnosed with lung cancer^6^. T cells were obtained from both peripheral blood and lung tumour tissues, including 44 distinct TCRs targeting four viral antigens (details provided in Supplementary Information). For each TCR, we computed its minimum Euclidean distance to those in the benchmark dataset. As shown in Supplementary Fig. 4b, 88.6% of antigen-specific TCRs exhibited a minimum distance greater than 0.1 from any TCR in the benchmark dataset, indicating limited overlap and unseen sequence distribution. To construct non-binding antigen–TCR pairs, we grouped the binding TCRs by antigen and randomly sampled an equal number of non-binding TCRs from the remaining pool. We used the five cross-validated models to predict binding probabilities for both positive and negative pairs, and averaged the results to obtain final binding scores. As shown in Fig. 5e, binding and non-binding pairs exhibited significantly different prediction score distributions (unpaired t-test, *p* < 0.0001). Furthermore, when we plotted the prediction scores of each clonotype by antigen (Fig. 5f), we found that 5 of the 44 TCRs recognized more than one antigen, suggesting cross-reactivity within the donor’s antigen-specific TCR repertoire. The 10 antigen–TCR pairs exhibiting cross-reactivity were highlighted in green. Notably, the cross-reactive TCRs exhibited consistently lower predicted binding scores compared to mono-specific counterparts. From a biological perspective, cross-reactive T cells enhance immune coverage by recognizing a broader array of peptides, often at the expense of antigen specificity^1^. Computationally, this phenomenon may be attributed to the model’s conservative prediction strategy in the absence of high-confidence, antigen-specific sequence features. This suggests that TridentTCR may prioritize precision over recall when encountering promiscuous TCRs—a behavior that aligns with the clinical requirement for high target specificity in T cell-based immunotherapy.

Collectively, these results demonstrate that TridentTCR robustly predicts antigen-specific interactions for previously unseen TCRs, and that its output scores can distinguish clonotypes with greater expansion potential from non-reactive T cells.

#### TridentTCR predicts the autoreactive T cell interactions for unseen arAg

To evaluate whether TridentTCR can predict arAg–TCR interactions for previously unseen peptides in disease-specific contexts, we curated a dataset from a study focusing on T1D^24^. This dataset included 14 T cell clonotypes reactive to the ZnT8_186-194_ (VAANIVLTV) epitope, an immunodominant target in T1D. Among them, six clonotypes were isolated from five T1D patients and eight from six healthy donors (details in Supplementary Information). For each TCR, we calculated its minimum Euclidean distance to all TCRs in the benchmark dataset. As shown in Supplementary Fig. 4c, 10 of these TCRs exhibited a minimum distance greater than 0.1, suggesting they are largely unseen by the model. We then applied the five cross-validated models to predict binding probabilities for all arAg–TCR pairs, averaging the results to obtain final prediction scores. As shown in Supplementary Fig. 6, when grouped by disease status, the average arAg–TCR prediction scores in the T1D group were slightly higher than those in the healthy group. However, this difference was not statistically significant (unpaired t-test). This is consistent with findings from Culina *et al.*, who reported that islet-reactive CD8^+^ T cells are broadly present in the general population and exhibit comparable functionality, including antigen-binding avidity^24^. Moreover, the overall distribution of arAg–TCR prediction scores was moderate, rather than skewed toward high-confidence predictions. This observation aligns with reports from the same study, which identified antigen-experienced T cells in both T1D patients and healthy individuals. These findings raise the possibility of cross-priming driven by homologous antigen KAANIVLTV, derived from the intestinal commensal *Bacteroides stercoris* (*B. stercoris*)^24,26^. Compared to VAANIVLTV, this antigen differs by a single N-terminal amino acid substitution. We computed the Euclidean distance between KAANIVLTV and all antigens in the benchmark dataset (Supplementary Fig. 4c). The minimum distance was 0.265, confirming that this arAg is novel and unseen by the model. We then predicted the binding scores of all 14 TCRs to the KAANIVLTV antigen. As shown in Fig. 5g, the predicted binding specificity to KAANIVLTV was significantly higher than that to VAANIVLTV, despite the fact that KAANIVLTV had never been seen by TridentTCR (unpaired t-test, *p* < 0.0001). This result aligns with prior observations that the *B. stercoris*–derived antigen displays stronger agonist potency than the ZnT8^186-194^ antigen, further supporting the notion that T cell cross-reactivity may play a role in the initiation of autoimmune responses^24^. Collectively, these findings demonstrate that TridentTCR can generalize to previously unseen arAgs and accurately identify TCR cross-reactivity driven by subtle sequence homology.

**Fig. 6.**
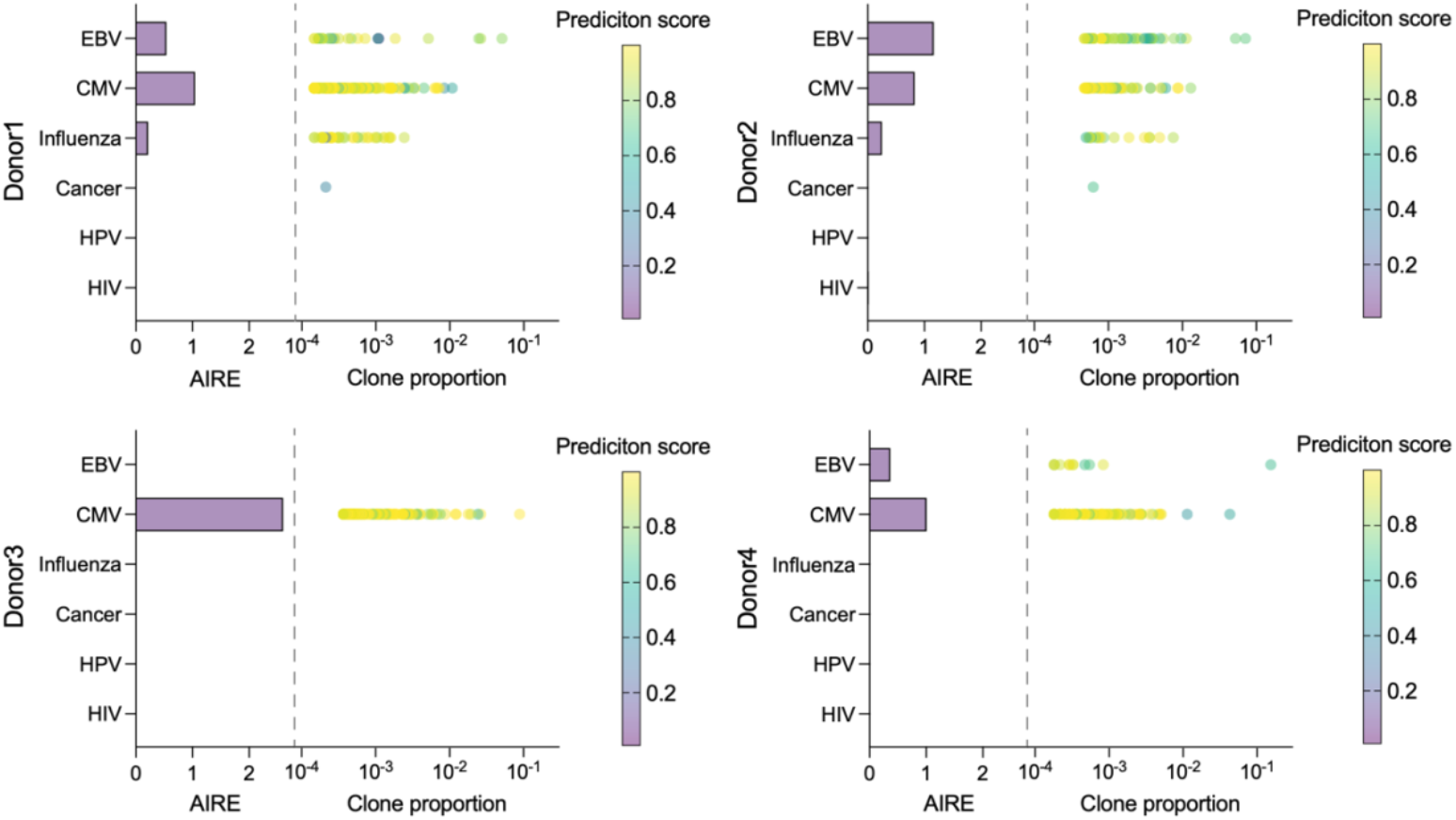
Integration of antigen–TCR interactions, clonotype frequencies, and repertoire diversity using the AIRE metric. For each donor, AIRE scores were calculated for six disease-associated antigens— EBV BZLF1_190-197_ (RAKFKQLL), CMV IE1_184-192_ (KLGGALQAK), Influenza A M1_58-66_ (GILGFVFTL), Cancer Tyrosinase_8-17_ (CLLWSFQTSA), HPV E7_12-20_ (MLDLQPETT), and HIV Gag_150-158_ (RTLNAWVKV)— based on their antigen-specific T cell repertoires. Bar height indicates the AIRE value for each antigen. Each dot represents a unique T cell clonotype, plotted by its clonal proportion (x-axis) and colored by its predicted binding probability to the corresponding antigen.

### Antigen-TCR interactions, clonotypes proportion and diversity infer antigenic immune response

Ideally, antigens presented on diseased cells should be recognized by CD8^+^ T cells to elicit immune responses that promote the eradication of these cells. However, accurately quantifying the magnitude of antigen-specific T cell responses remains challenging due to multiple contributing factors. First, the probability that a given T cell clonotype can recognize a specific antigen determines its functional relevance, forming the basis of antigen-specific immune activation. Second, the frequency of antigen-reactive T cell clones also plays a critical role. For instance, in type 1 diabetes, increased frequencies of islet-reactive T cells in the pancreas distinguish patients from healthy donors, suggesting that clonal abundance provides complementary predictive value. Third, the diversity of the antigen-specific T cell repertoire contributes to the heterogeneity of T cell activation states and may partially explain inter-patient variability in response to immunotherapy^27,28^. Together, antigen–TCR interactions, clonotype frequencies, and TCR repertoire diversity shape the magnitude and quality of the immune response to specific antigens. A study in patients with metastatic castration-resistant prostate cancer and melanoma demonstrated that CTLA-4 blockade induces T cell repertoire diversification. Moreover, the maintenance of certain high-frequency, antigen-specific T cell clones was associated with more favorable clinical outcomes^29^.

To quantify this effect, we introduced a metric AIRE, inspired by the concept of information entropy. AIRE is computed as the weighted sum of the contributions of all antigen-specific T cell clonotypes, where each weight is defined as the product of the antigen–TCR recognition probability and the corresponding clonotype frequency (see Supplementary Information for details). AIRE simultaneously accounts for the clonal diversity, expansion levels, and functional antigen recognition efficacy of antigen-specific T cell clones. It thus serves as a quantitative indicator of the immune memory landscape. A high AIRE value suggests a more diverse and redundant antigen-specific T cell response, which may lead to a stronger overall immune activation. Under homeostatic conditions, elevated AIRE scores may reflect enhanced immune surveillance capacity.

We investigated the association between AIRE values and disease status across a cohort of four donors from the 10x Genomics dataset, focusing on their antigen-specific T cell repertoires. We analyzed responses to six disease-associated antigens: EBV BZLF1_190-197_ (RAKFKQLL), CMV IE1_184-192_ (KLGGALQAK), Influenza A M1_58-66_ (GILGFVFTL), Cancer Tyrosinase_8-17_ (CLLWSFQTSA), HPV E7_12-20_ (MLDLQPETT), and HIV Gag_150-158_ (RTLNAWVKV). We first examined donor 4, who served as a seronegative control and was reported to be seronegative for CMV, EBV, and HIV. Although the disease status for Influenza, cancer, and HPV was not reported, no antigen-specific T cell clonotypes were detected for these antigens in donor 4. These findings suggest that the AIRE values of donor 4 (AIRE_EBV_ = 0.368, AIRE_CMV_ = 1.01) may reflect baseline immune surveillance without detectable antigen-specific responses or evidence of prior infection. We further analyzed donors 1, 2, and 3, who were seropositive for one or more antigens, and found that elevated AIRE values were positively associated with prior antigen exposure. Among donors 1, 2, and 4, AIRE values were markedly higher for lifelong latent viruses such as EBV and CMV, compared to non-exposed or infrequently encountered antigens such as HPV and HIV. This observation is potentially explained by the persistence of high-frequency memory CD8^+^ T cell clonotypes in peripheral blood following previous infections. Moreover, donor 1 (AIRE_EBV_ = 0.542, AIRE_Influenza_ = 0.214) and donor 2 (AIRE_EBV_ = 1.16, AIRE_Influenza_ = 0.251) exhibited significantly higher AIRE scores for EBV and Influenza compared to donor 4. These responses were accompanied by more abundant and higher-recognition probability antigen-specific clonotypes, consistent with their reported seropositive status for these pathogens. Interestingly, as shown in Supplementary Fig. 7, the distribution of clonotype contributions to the Influenza response differed markedly between donors 1 and 2 (*p* = 0.0003). Donor 2’s repertoire was dominated by a few highly expanded and high-recognition probability clonotypes, while donor 1 exhibited a broader but less expanded set of clones with lower contributions. Both configurations suggest effective but distinct immune strategies. Additionally, donor 3 displayed the highest AIRE value for CMV (AIRE_CMV_ = 2.60), with a diverse set of high recognition probability antigen-specific T cell clonotypes, consistent with their known seropositive status. Collectively, by quantifying the diversity–efficacy trade-off across clonotypes, AIRE integrates breadth (clonal diversity), depth (clonal expansion), and quality (predicted binding probability) into a unified metric of immunological memory. This measure captures the immunological imprint of past infections and may serve as a proxy for immune reserve. Its observed correlation with serostatus highlights its potential utility in assessing vaccine responsiveness and forecasting clinical outcomes.

## Discussion

Developing *in silico* methods to predict antigen–TCR interactions provides a cost-effective and efficient pathway for enhancing T cell-based immunotherapy. Despite substantial advances, accurately characterizing antigen recognition by TCRs remains challenging due to individual TCR repertoire diversity and autoimmune-related toxicities arising from T cell cross-reactivity. To address this, we developed TridentTCR, a computational framework designed to refine TCR specificity predictions systematically. Unlike conventional methods, which typically adopt binary classification to predict antigen-TCR binding, TridentTCR uniquely incorporates a trinary classification task. Specifically, its binary mode differentiates binding from non-binding antigen-TCR interactions, while the trinary mode further distinguishes TCR interactions involving general antigens from arAgs.

Importantly, our definition of arAgs is broader than traditional self-antigens, encompassing any antigen associated with autoimmune pathogenesis through diverse mechanisms. For instance, the EBV antigen EBNA-1 exemplifies such complexity, as EBV infection significantly elevates multiple sclerosis risk—potentially via molecular mimicry, triggering autoreactive T cell responses ^30,31^. Our application of TridentTCR to three independent clinical datasets demonstrated robust generalization to previously unseen TCR sequences and antigens. In a disease-specific investigation, we identified that ZnT8_186-194_-specific TCRs preferentially recognize *B. stercoris* antigen, suggesting T cell cross-reactivity as a potential contributor to T1D pathogenesis. Furthermore, leveraging TridentTCR’s predictive capability on single-cell immune profiling data, we showed that our AIRE metric has prognostic potential and could serve as a predictive marker for immunotherapy response.

Nevertheless, this study faces limitations primarily related to dataset biases inherent in publicly available antigen–TCR interactions. Current datasets disproportionately represent TCR sequences relative to antigen diversity, due largely to experimental constraints associated with pMHC-I tetramer generation. Tetramer-based assays, although highly specific, are costly, especially for rare MHC alleles lacking commercial availability, further limiting antigen diversity. Additionally, comprehensive annotation of arAgs that induce cross-reactivity remains incomplete.

In future work, continued accumulation of balanced antigen–TCR and arAg–TCR interaction datasets will likely improve TridentTCR’s specificity and predictive performance, fostering a deeper understanding of TCR recognition mechanisms.

In summary, TridentTCR represents a unified and clinically oriented computational framework capable of systematically distinguishing TCR specificity toward general and autoimmune-related antigens. We anticipate TridentTCR will contribute to immunogenomics research, informing safer, more precise immunotherapy strategies, and ultimately improving patient outcomes.

## Methods

### Datasets

#### Benchmark dataset curation

The benchmark dataset used for training TridentTCR was curated from publicly available databases, including McPAS-TCR, IEDB, and VDJdb (collection dates and URLs provided in Supplementary Information)^19-21^. We included only human TCRβ CDR3 sequences restricted to human leukocyte antigen class I (HLA-I) alleles. Records were filtered according to the following criteria: (1) Retained TCRβ CDR3 sequences ranging from 10 to 20 amino acids, and antigen sequences ranging from 8 to 15 amino acids; (2) Removed sequences containing lowercase letters, non-standard amino acids, or duplicate antigen–TCR sequence pairs; (3) Applied iSMART, a high-performance TCR clustering tool based on pairwise local alignment algorithms, to cluster TCR sequences into antigen-specific groups, and removed singleton TCRβ CDR3 sequences that were not assigned to any clusters^9^. The final curated benchmark dataset contained 41,732 antigen–TCR binding pairs. Since available databases provide very few experimentally verified non-binding antigen–TCR pairs, insufficient for robust model training, we generated negative interaction pairs computationally. For each antigen, we constructed an equal number of non-binding pairs by randomly pairing the antigen with TCR sequences that have no known binding interactions with that antigen.

#### External validation datasets curation

To rigorously evaluate model performance and assess its clinical applicability, we assembled three independent external validation datasets obtained from publicly available sources, including 10x Genomics, Lu *et al.*, and Culina *et al* ^6,23,24^. These datasets comprised antigen–TCR binding pairs with associated metadata: 612 pairs from 10x Genomics, 49 from Lu *et al.*, and 16 from Culina *et al.* (Supplementary Information). Due to the limited availability and small scale of experimentally verified antigen–TCR pairs typically encountered in clinical trial scenarios, we did not apply the iSMART clustering-based filtering step used for benchmark dataset curation. This approach better reflects real-world clinical conditions, ensuring that the validation accurately assesses the model’s generalizability in scenarios characterized by high sequence diversity and limited data volume.

### Defining arAgs

The arAgs were broadly defined as peptides associated with autoimmune pathogenesis through various mechanisms, not limited to classical self-antigens. We extracted candidate arAgs from two databases: IEDB and McPAS-TCR ^19,21^. From IEDB, we selected peptides derived from human subjects diagnosed with autoimmune diseases and restricted to HLA-I alleles. From McPAS-TCR, we included peptides annotated under the “autoimmune” disease category and derived from human subjects. Peptides from both sources were merged and further filtered to retain only those ranging from 8 to 15 amino acids, and to exclude sequences containing lowercase letters or non-standard amino acid characters. In total, 767 arAgs were obtained from IEDB and 19 from McPAS-TCR.

### Embedding antigen and TCR sequences using ESM-2

Antigen and TCRβ CDR3 sequences were encoded using ESM-2, a RoBERTa-based protein language model pretrained with an unsupervised masked language modeling objective, which generates high-dimensional embeddings that capture contextual and evolutionary information of the sequences ^17^. We employed the medium-sized version of ESM-2 (33-layer transformer, 650 million parameters), pretrained on billions of protein sequences, providing a balanced trade-off between computational efficiency and representational power. The embedding procedure was conducted as follows: (1) Each input sequence was tokenized into discrete tokens. (2) Tokens were fed into ESM-2 to obtain per-residue representations. (3) Per-residue embeddings were averaged to generate 1,280-dimensional sequence-level embeddings. The resulting embeddings for an antigen *a* ∈ 𝒜 and a TCR sequence *t* ∈ 𝒯 were denoted as 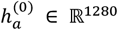 and 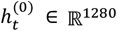, respectively. Although we currently focused only on the CDR3 region, which predominantly determines antigen specificity, future implementations may also incorporate CDR1 and CDR2 regions. This extension is feasible, considering that ESM-2 accepts sequences up to 1,022 amino acids in length.

### Latent neighbourhoods construction

We construct latent edges based on node features projected into a latent space, aiming to capture node pairs that are not directly connected in the original graph but are close in feature space. These latent edges augment the structural information of the original graph by encoding feature-level similarity. For each antigen *a* ∈ 𝒜 and TCR sequence *t* ∈ 𝒯, the initial feature vectors are denoted as 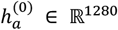 and 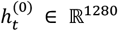, respectively. A dimensionality reduction function *f*: ℝ^1280^ → ℝ^*k*^ is applied to project these features into a latent space, yielding embeddings 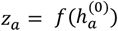 and 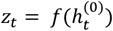. The latent similarity edges for antigens and TCRs are defined as follows:

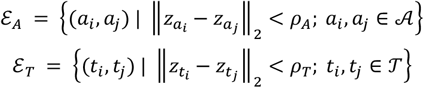

Here, *ρ*_*A*_ and *ρ*_*T*_ are not fixed hyperparameters, but are instead adaptively determined to ensure that the latent neighborhoods match the structural density of the original graph. We gradually increase the radius *ρ*_*A*_ and *ρ*_*T*_ for antigens and TCRs until the average number of neighbors in the latent space meets or exceeds that in the original graph. And ‖ · ‖_2_ denotes the Euclidean distance in the latent space. Compared with the observed antigen–TCR interaction edges ℰ_*inter*_, the latent similarity edges ℰ_*A*_and ℰ_*T*_may connect node pairs that are structurally distant in the original graph but are semantically close in the latent space. Biologically, such edges capture shared reactivity patterns—e.g., antigens that tend to be recognized by similar TCRs, or TCRs that target similar antigens.

Additionally, for each latent edge (*v*_*i*_, *v*_*j*_) ∈ ℰ_*A*_∪ ℰ_*T*_, we further assign a discrete geometric relationship label using a relational operator *τ*, which maps the relative positions of the two nodes in latent space to a finite relation set ℛ:

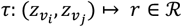

In this study, ℛ = {upper-left,upper-right,lower-left,lower-right}, where *r* indicates the quadrant position of node *v*_*j*_ with respect to node *v*_*i*_ in a 2D Euclidean space.

### Multi-view geometric graph learning

To jointly capture observed antigen–TCR cross-type interactions and latent intra-type similarities of TCRs or antigens, we construct a unified graph 𝒢 = (𝒱, ℰ), where 𝒱 = 𝒯 ∪ 𝒜 denotes the node set containing TCRs 𝒯 and antigens 𝒜, including both general antigens and arAgs. Each node is initialized with its corresponding embedding, denoted as 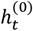 for *t* ∈ 𝒯 and 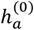 for *a* ∈ 𝒜. The edge set ℰ = {ℰ_*inter*_, ℰ_*A*_, ℰ_*T*_} includes three relation types: observed antigen–TCR interactions ℰ_*inter*_, antigen latent similarity edges ℰ_*A*_, and TCR latent similarity edges ℰ_*T*_, both ℰ_*A*_and ℰ_*T*_were constructed in the latent space via pairwise Euclidean proximity.

We propose a bi-level multi-view geometric graph learning framework to integrate the two views. At the low-level, message passing is performed independently on each view. Specifically, we apply GNN modules on (1) the interaction view to capture cross-type binding information, and (2) the latent similarity views to capture intra-type geometric structure. For each node *v* ∈ 𝒱 at layer *l*, its interaction-view feature is updated as:

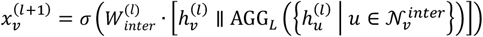

where *AGG* (·) is a permutation-invariant mean aggregator, 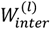 is a trainable weight matrix, and ∥ denotes feature concatenation.

In the latent views, message passing is conditioned on geometric relations *r* ∈ ℛ, determined by the quadrant direction from *z*_*v*_ to *z*_*u*_ in the latent space. The updated feature under each relation *r* is:

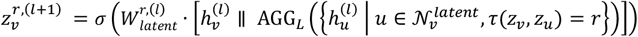

At the high-level, the outputs from interaction and latent views are fused using another aggregation function *AGG*_*H*_(·) to generate the unified representation:

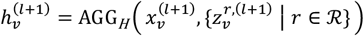

This fusion integrates biochemical interaction evidence and latent structural similarity, enabling the model to generalize across both seen and unseen antigen–TCR pairs.

### Learning TCR binding specificity for antigens and arAgs

The multi-view geometric graph learning framework generates embeddings for TCRs and antigens in the form of numerical vectors. To predict antigen–TCR binding specificity, the embeddings of antigen–TCR pairs are concatenated and subsequently passed into a MLP classifier. The integrated embeddings are first processed through a fully connected layer with 512 neurons followed by ReLU activation, and then a second fully connected layer with 256 neurons and ReLU activation. For the trinary classification setting, the final output layer contains three neurons activated by a softmax function, producing a vector of three probabilities representing the categories: general antigen–TCR binding, arAg–TCR binding, and no interaction. In the binary classification scenario, the final layer consists of a single neuron activated by a sigmoid function, outputting the binding probability between the given antigen–TCR pair.

Formally, for each antigen *a* ∈ 𝒜 and TCR sequence *t* ∈ 𝒯, we denote the predicted probability as 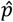, and the corresponding ground-truth interaction as *p*. In the binary setting, *p* ∈ {0, 1}, representing non-binding and binding interactions, respectively. In the trinary setting, we define *p* ∈ {0, 1, 2}, further differentiating binding interactions into general antigen-binding (label 1) and arAg-binding (label 2).

### Hybrid loss function

To focus model training on challenging, uncertain, or underrepresented cases, we design a hybrid loss function combining the cross-entropy loss and focal Tversky loss:

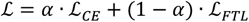

where *α* is a scalar hyperparameter balancing the contributions of the two loss terms. The cross-entropy loss ℒ_*CE*_is computed by 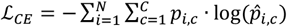, *N* is the total number of antigen– TCR pairs, *C* denotes the number of interaction classes (binary: *C* = 2; trinary: *C* = 3), *p*_*i,c*_is the one-hot encoding of the true label, and 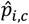 is the predicted probability for the *i*-th sample belonging to class *c*.

The focal Tversky loss *ℒ*_*FTL*_ is defined by *ℒ*_*FTL*_ = (1 − *T*_*C*_)^*γ*^, where *γ* is a focusing parameter (set to 2 by default), and 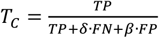 is the Tversky index calculated over all classes, with *δ* controlling penalties for false negatives (default 0.7), and *β* controlling penalties for false positives (default 0.3). The TP (true positives), FP (false positives), and FN (false negatives) are computed as: 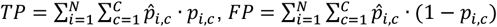, and 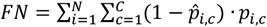. The hybrid loss thus emphasizes challenging classification scenarios and improves the model’s robustness and discriminative ability.

The TridentTCR model was trained for 1,000 epochs using full-batch gradient descent optimized with Adam (learning rate =0.001). To reduce memory consumption and accelerate training, automatic mixed-precision training was employed.

## Acknowledgements

This study was supported by the National Natural Science Foundation of China (Grants No. 62433016 and 624B2112, awarded to X.S., R.N., respectively). We gratefully acknowledge these funding agencies for their support.

## Author contributions

X.S. conceived and supervised the study. R.N., X.K., and J.Z. conducted the computational analysis and tested the methods. R.N. wrote the initial draft of the manuscript. Y.L., and X.S. critically reviewed and edited the manuscript.

## Competing interests

The authors declare no competing interests.

## Notes

### Competing Interest Statement

The authors have declared no competing interest.

