## Supplementary Information for "TridentTCR: Predicting T Cell Receptor Specificity for Antigens and Autoimmune-Related Targets via a Topology-Aware Graph and Large Language Model"

### 1. Data summary

**Supplementary Tab. 1 Summary of datasets used for model training and validation.**

| Cohort | Ref. | Type | Number | Role | Methodology | Comments |
|---|---|---|---|---|---|---|
| IEDB | 1 | Antigen-TCR binding pairs | 110,384 | Training | Database-curated antigen specificity | 5-fold cross-validation |
| VDJdb | 1 | Antigen-TCR binding pairs | 4,151 (Confidence score =1); 1,662 (Confidence score = 2); 783 (Confidence score = 3) | Training | Database-curated antigen specificity | 5-fold cross-validation |
| McPAS-TCR | 2 | Antigen-TCR binding pairs | 9,696 | Training | Database-curated antigen specificity | 5-fold cross-validation |
| IEDB | 1 | Autoimmune-related antigens (arAgs) | 767 | Training | Metadata annotation | Identified arAgs based on database metadata |
| McPAS-TCR | 2 | arAgs | 19 | Training | Metadata annotation | Identified arAgs based on database metadata |
| 10x Genomics | 3 | Single-cell antigen specificity and TCR repertoire (CDR3 sequences, clone proportion, donor metadata) | 80,110 T cell clonotypes (44 pMHC multimers) | Testing | Single-cell immune profiling (10x Genomics Chromium) | External validation for generalization performance and antigenic immune response entropy (AIRE) calculation |
| Lu *et al.* | 4 | Antigen–TCR binding pairs (virus-specific) | 49 | Testing | Single-cell TCR sequencing after antigen-specific *in vitro* expansion | External validation (virus-infected donor) |
| Culina *et al.* | 5 | T1D antigen specific TCR sequences and recognition specificity to B. stercoris mimotope | 16 | Testing | Multimer staining assays | External validation (T1D and healthy donors) |

The benchmark dataset was curated from three well-known sources: McPAS-TCR, IEDB, and VDJdb. The IEDB dataset was downloaded from https://www.iedb.org/ on October 11, 2024. VDJdb was obtained from https://vdjdb.cdr3.net on the same date, and McPAS-TCR was retrieved from https://friedmanlab.weizmann.ac.il/McPAS-TCR/ on June 14, 2024.

### 2. TCR Similarity Analysis and Minimum Distance Computation

To evaluate the robustness of our model under varying levels of TCR similarity between training and test sets, we partitioned the benchmark dataset into 80% training and 20% test data. The similarity between test and training TCRs was quantified by computing euclidean distances in a reduced feature space.

Specifically, we first encoded all TCR sequences using the pre-trained ESM-2 model, resulting in 1280-dimensional embeddings. These embeddings were concatenated and projected into a 30-dimensional space using uniform manifold approximation and projection (UMAP) for distance-based comparison.

Let the test set contain $m$ TCRs and the training set contain $n$ TCRs. For each test TCR $\mathrm{tcr_i}$, we computed its euclidean distance to every training TCR $\mathrm{TCR}_j$, denoted as $dij \in R$. For each test TCR, only the minimum distance was retained:

$$D_i = \min_{j \in [1,n]} d_{ij}, \quad \text{for } i = 1,2,\ldots,m$$

This results in a distance matrix of size $m \times n$, with an additional column recording the minimum distance $D_i$ for each test TCR. Among all minimum distances $\{D_i\}_{i=1}^{m}$, the smallest observed value was 0.0005 and the largest was 0.8538. This distribution exhibits a long-tailed shape, indicating a wide spectrum of similarity between test and training TCRs. The full distribution of minimum distances is shown in **Supplementary Fig. 1**, where a histogram illustrates the number of test TCRs at each distance interval, along with a fitted Gaussian curve. Notably, 90% of test TCRs have a minimum distance less than 0.172, suggesting that most test samples fall within a moderate similarity range relative to the training set.

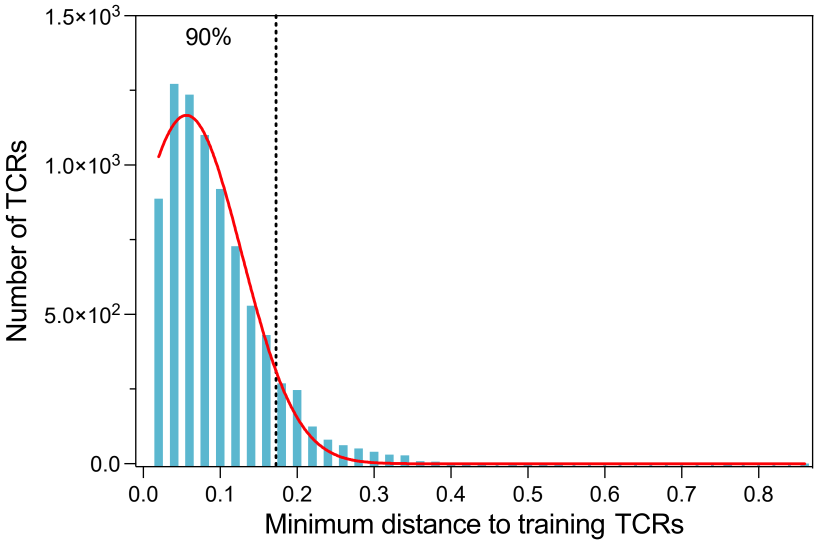


**Supplementary Fig. 1 Distribution of test TCR similarity to training TCRs.** The histogram shows the number of test TCRs within each 0.02-wide distance bin, with the x-axis representing the minimum distance to training TCRs and the y-axis indicating the number of TCRs. A Gaussian curve was fitted to the distribution (red line). The results reveal a long-tailed distribution of TCR distances, reflecting varying levels of similarity between test and training sequences.

### 3. Visualization and Evaluation of TCR Embedding Using Different Encoding Methods

To evaluate the representational quality of different TCR encoding methods, we compared four approaches: (1) the ESM-2 pretrained language model used in TridentTCR, (2) the biologically informed substitution matrix BLOSUM50, (3) the convolutional neural network (CNN)-based encoder used in HeteroTCR, and (4) the autoencoder architecture adopted by pMTnet[6,7,8]. 8,506 unique TCRβ CDR3 sequences were obtained from VDJdb and embedded using each respective method[1]. The resulting high-dimensional embeddings were reduced to two dimensions using UMAP, and the recognized epitope species served as labels for clustering evaluation.

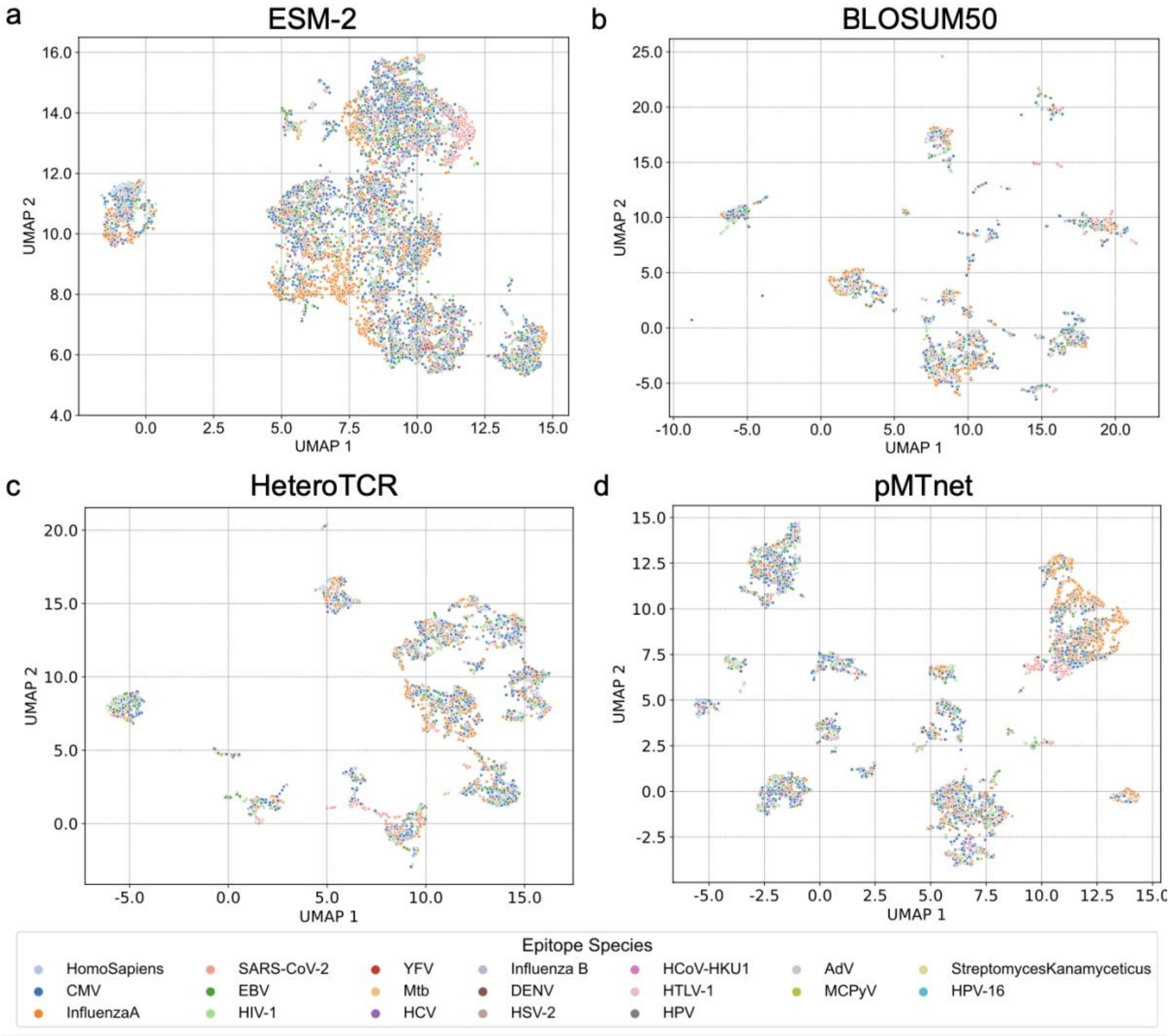


**Supplementary Fig. 2 UMAP projections of TCR embeddings from different encoding methods.** UMAP visualization of TCR embeddings derived from four encoding methods: **a**, ESM-2; **b**, BLOSUM50; **c**, HeteroTCR's pre-trained CNN encoder; and **d**, pMTnet's pre-trained autoencoder. Each point represents a TCR from the VDJdb dataset and is colored by its corresponding epitope species.

As shown in **Supplementary Fig. 2**, TCRs encoded by BLOSUM50 formed scattered micro-clusters without clear semantic grouping. Many clusters were internally color-mixed, indicating poor aggregation of TCRs recognizing the same epitope species. This reflects the biochemical substitution pattern captured by BLOSUM50, which is effective at low-level sequence similarity but fails to capture high-level immune semantics. The pre-trained CNN encoder from HeteroTCR and the autoencoder from pMTnet produced more compact distributions than BLOSUM50. Particularly, pMTnet demonstrated improved intra-class aggregation, with TCRs recognizing the same epitope tending to cluster together. However, overlapping regions between different classes were still observed in both models, suggesting limited class separation and semantic leakage.

In contrast, the pre-trained protein language model ESM-2 yielded a markedly better latent structure. TCRs recognizing the same epitope species were grouped into visually coherent clusters with consistent color, and boundaries between epitope classes appeared smoother and more biologically plausible. Notably, distinct regions were dominated by specific epitope species—for example, CMV-specific TCRs formed a compact cluster in the upper left, and Influenza A–specific TCRs aggregated on the right—suggesting that ESM-2 embeddings capture both intra-class consistency and inter-class separability.

We then assessed cluster separability using the **Davies–Bouldin (DB) index**, which quantifies intra-cluster compactness and inter-cluster separation. A lower DB index indicates better class-wise separation in the latent space. Let $C_k$ denote the set of TCR embeddings belonging to epitope species $k$, and $\mu_k$ its centroid. Then the DB index is computed as:

$$\text{DB} = \frac{1}{K}\sum_{i=1}^{K} \max_{j\neq i}\left(\frac{\sigma_i + \sigma_j}{\|\mu_i - \mu_j\|}\right)$$

where $\sigma_k$ is the average distance from each point in $C_k$ to its centroid $\mu_k$, and $K$ is the number of epitope species.

To further assess whether embeddings preserve **fine-grained feature within each epitope species**, we computed two intra-class metrics: (1) the **mean intra-class pairwise distance** (the average euclidean distance between all TCRs within the same epitope species), and (2) the **mean intra-class variance**, computed as the trace of the covariance matrix of the embedded TCRs in each class. Formally, for class $C_k = \{x_1, x_2, \ldots, x_{n_k}\}$, we define:

$$\text{Intra-class mean distance}_k = \frac{2}{n_k(n_k - 1)}\sum i < j \,\|x_i - x_j\|$$

$$\text{Intra-class variance}_k = \text{Tr}\big(\text{Cov}(C_k)\big)$$

where $\text{Cov}(C_k)$ is the sample covariance matrix of $C_k$, and $Tr$ denotes the trace operator.

Averaged results across all epitope species are reported in **Supplementary Tab. 2**, including the DB index, mean intra-class distance, and intra-class variance for each encoding method. Detailed per-epitope statistics are presented in **Supplementary Tab. 3** and **Supplementary Tab. 4**.

In the overall clustering performance assessment (**Supplementary Tab. 2**), ESM-2 achieved the best results across all three metrics: the lowest Davies–Bouldin (DB) index, the lowest intra-class mean distance, and the lowest intra-class variance. Furthermore, in the per-epitope intra-class analysis (**Supplementary Tab. 3 and Tab. 4**), ESM-2 consistently exhibited the lowest intra-class mean distance and variance across nearly all epitope species, with the sole exception of AdV.

**Supplementary Tab.2 Overall evaluation of TCR embedding quality across different encoding strategies.**

| Matrix | TridentTCR (ESM-2) | BLOSUM50 | HeteroTCR (CNN) | pMTnet (Autoencoder) |
|---|---|---|---|---|
| DB index | **19.8** | 69.0 | 27.7 | 31.1 |
| Average intra-class distance | **4.92** | 11.1 | 9.04 | 8.68 |
| Average intra-class variance | **8.47** | 40.8 | 26.2 | 24.9 |

**Supplementary Tab.3 Per-epitope species intra-class distance for different TCR encoding strategies.**

| Epitope Species | TridentTCR (ESM-2) | BLOSUM50 | HeteroTCR (CNN) | pMTnet (Autoencoder) |
|---|---|---|---|---|
| AdV | 2.96 | **0.009** | 0.324 | 0.206 |
| CMV | **5.55** | 11.9 | 9.48 | 9.35 |
| DENV | **5.08** | 10.2 | 7.79 | 9.6 |
| EBV | **5.56** | 13.2 | 10.1 | 9.18 |
| HCV | **5.34** | 10.7 | 8.71 | 9.06 |
| HIV-1 | **5.73** | 12.5 | 10.2 | 9.24 |
| HPV | **2.6** | 17.8 | 20.1 | 9.95 |
| HSV-2 | **2.81** | 9.33 | 5.78 | 8.65 |
| HTLV-1 | **5.2** | 10.3 | 8.92 | 7.76 |
| HomoSapiens | **6.22** | 12.2 | 9.31 | 9.9 |
| Influenza B | **6.46** | 11.5 | 8.33 | 10.3 |
| InfluenzaA | **5.42** | 11.8 | 8.58 | 9.37 |
| Mtb | **5.26** | 11.8 | 9.82 | 9.71 |

| | | | | |
|---|---|---|---|---|
| SARS-CoV-2 | **5.1** | 12.7 | 9.08 | 8.74 |
| YFV | **4.63** | 11.9 | 9.05 | 9.27 |

**Supplementary Tab.4 Per-epitope species intra-class variance for different TCR encoding strategies.**

| Epitope Species | TridentTCR (ESM-2) | BLOSUM50 | HeteroTCR (CNN) | pMTnet (Autoencoder) |
|---|---|---|---|---|
| AdV | 2.33 | **1.74E-05** | 0.0259 | 0.00971 |
| CMV | **10.4** | 45.9 | 29.8 | 27.6 |
| DENV | **8.78** | 34.6 | 21.5 | 29.2 |
| EBV | **10.2** | 56.4 | 33.2 | 26.6 |
| HCV | **9.85** | 36.9 | 24.6 | 26.9 |
| HIV-1 | **11.2** | 50.2 | 33.9 | 27.2 |
| HPV | **0.846** | 39.7 | 50.3 | 12.4 |
| HSV-2 | **2.16** | 24.9 | 9.43 | 24.8 |
| HTLV-1 | **8.93** | 36.2 | 26.6 | 22.4 |
| HomoSapiens | **13** | 47.9 | 28.5 | 31 |
| Influenza B | **13.3** | 41.5 | 22.7 | 33.7 |
| InfluenzaA | **9.97** | 46.4 | 25.3 | 29.4 |
| Mtb | **9.1** | 44.7 | 30.6 | 29.2 |
| SARS-CoV-2 | **8.92** | 51.9 | 27.1 | 24.9 |
| YFV | **8.19** | 55.2 | 30.1 | 28.4 |

### 4. Ablation Study to Quantify the Contribution of Sequence and Network Representations

To evaluate the respective contributions of sequence-derived and topology-based information to TridentTCR's performance, we conducted ablation experiments on the three-class dataset (antigen–TCR, arAg–TCR, and non-interacting pairs).

We designed two ablated variants of the original TridentTCR model:

- A feature-only variant, in which ESM-2 embeddings were directly fed into a multilayer perceptron (MLP) classifier, omitting graph-based learning entirely.
- A network-only variant, in which BLOSUM50 matrices were used to represent TCR sequences (to provide minimal sequence-level information), and the embeddings were passed through graph neural networks (GNNs) and MLP classifier as described previously.

To statistically validate these observations, we compared the performance of TridentTCR with both ablated models using a paired t-test across the six evaluation metrics. As shown in **Supplementary Fig. 3**, TridentTCR significantly outperformed the ESM-2 + MLP variant ($p$ = 0.0002) and the GNN + MLP variant ($*** \ p$ < 0.0001).

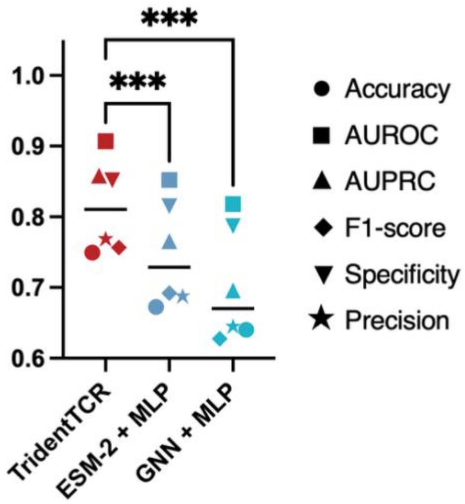


**Supplementary Fig. 3 Statistical comparison of TridentTCR with ablated model variants across six evaluation metrics.** Performance of TridentTCR, the feature-only variant (ESM-2 + MLP), and the network-based variant (GNN + MLP) was assessed using six metrics: accuracy, AUROC, AUPRC, F1-score, specificity, and precision. Each point represents the mean score for one metric across five folds. TridentTCR outperformed both ablated models significantly (paired t-test, $p$ = 0.0002 vs. ESM-2 + MLP; *** $p$ < 0.0001 vs. GNN + MLP).

### 5. Data curation and analysis of 10x Genomics Cohort

This dataset utilizes the 10x Genomics Chromium Single Cell Immune Profiling platform with Feature Barcode technology to directly profile antigen binding at the single-cell level [3]. It assesses the binding specificities of over 150,000 $CD8^+$ T cells from four human donors against a panel of 44 distinct peptide–MHC (pMHC) multimers (publicly available at 10x Genomics: https://www.10xgenomics.com/datasets). In this study, we utilized the standardized datasets provided by 10x Genomics for all four donors to investigate three key questions: (1) the inter-individual diversity of antigen-specific TCR repertoires; (2) whether the binding scores predicted by TridentTCR can qualitatively distinguish clonally expanded antigen-binding versus non-binding T cell clonotypes; (3) how entropy-based metrics can be applied to quantify the immune response strength of each donor to specific antigens. Accordingly, we curated the following data for each donor: (1) clone proportion and TCRβ CDR3 sequence of each T cell clonotype; (2) antigen specificity for each clonotype; and (3) donor metadata.

To identify antigen-specific T cell clonotypes for each donor, we performed the following steps. The binding specificity between T cells and pMHC multimers was assessed using unique molecular identifier (UMI) counts as a binding indicator. Clonotypes with a UMI count greater than 10 for at least one antigen-specific tetramer were retained. We then selected the top 1% of clonotypes based on their clone proportions. These clonotypes were designated as antigen-specific, and the corresponding antigen–TCR binding pairs were used as positive samples.

To assess whether the predicted binding scores could differentiate between expanded antigen-binding and non-binding clonotypes, we constructed a donor-specific negative set matched in size to the positive samples. Specifically, we first excluded the top 1% of clonotypes based on their clone proportions. To ensure that no antigen-binding signal was present, we removed all clonotypes with nonzero UMI counts across any tetramer. From the remaining pool, we randomly sampled an equal number of clonotypes to match the positive set for each donor. Finally, we grouped the positive antigen–TCR binding pairs by antigen and sampled an equal number of negative TCRs per antigen to generate corresponding non-binding antigen–TCR pairs.

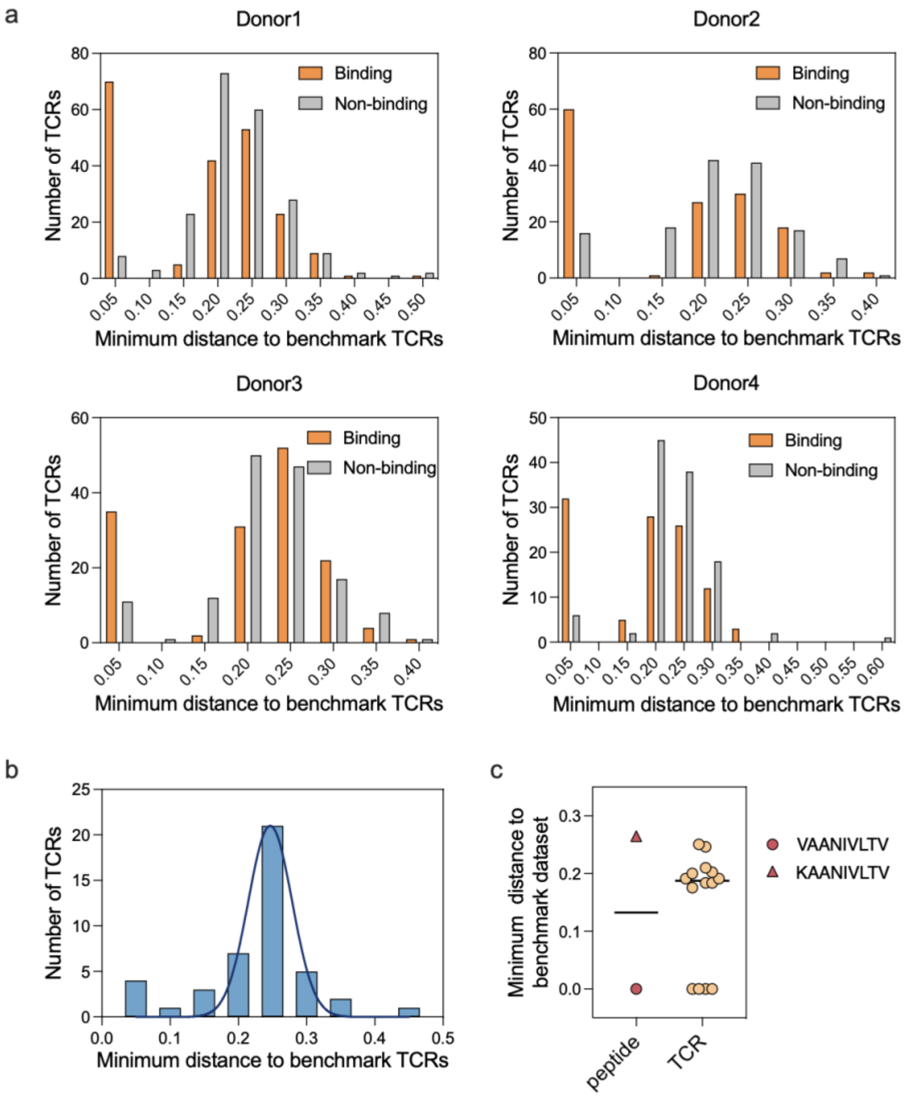


**Supplementary Fig. 4 Distance distribution of three independent evaluation datasets to the benchmark dataset.** **a**, Distribution of minimum euclidean distances between TCRs from the 10x Genomics dataset and TCRs in the benchmark dataset across four donors. TCRs are colored by antigen binding specificity. **b**, Distribution of minimum distances between 44 viral antigen–reactive TCRs and benchmark TCRs in a lung cancer patient with prior viral exposures. A Gaussian curve is overlaid for visualization. **c**, Minimum distances between peptides and benchmark peptides (left), and between ZnT8-reactive TCRs and benchmark TCRs (right). Each point represents one peptide or TCR; horizontal bars denote median values.

Subsequently, we used the pre-trained ESM-2 model to generate 1,280-dimensional embeddings for both binding and non-binding TCRs. We then concatenated these embeddings with those from the benchmark dataset and projected the combined data into a 30-dimensional space using UMAP for distance-based visualization and analysis. Finally, we computed the minimum Euclidean distance between each TCR from the 10x Genomics dataset and every TCR in the benchmark set. The full distribution of these minimum distances is presented in **Supplementary Fig. 4a**. Overall, more than 60% of antigen-specific TCRs from each donor had a minimum distance greater than 0.1, indicating that most of these TCRs are unseen by the model. In contrast, the negative TCRs were, on average, more distant from the benchmark set than the antigen-specific ones.

We then applied the five cross-validated TridentTCR models to predict the binding probabilities of both known binding and non-binding antigen–TCR pairs in an independent dataset. For each pair, the predicted scores from all five models were averaged to obtain the final binding probability. To further assess the statistical separation between groups, we visualized the prediction score distributions using violin plots for each donor (**Supplementary Fig. 5**).

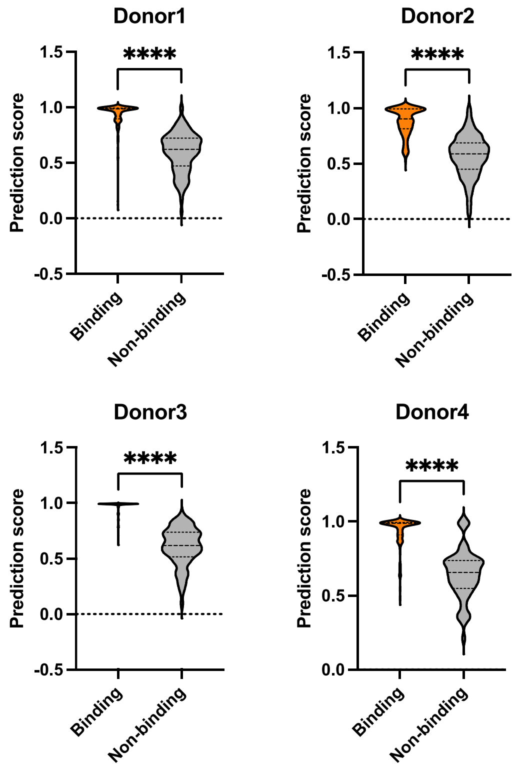


**Supplementary Fig. 5 Distributions of predicted binding scores for binding and non-binding clonotypes across four donors in 10x Genomics.** Binding probabilities were predicted using five cross-validated TridentTCR models and averaged to obtain final prediction scores. Violin plots show the distributions of binding scores for true binding (orange) and non-binding (gray) TCR–antigen pairs from four donors. In each case, antigen-specific clonotypes exhibited significantly higher prediction scores than non-binding clonotypes (unpaired t-test, $**** \ p$ < 0.0001), indicating robust discrimination by the model. The boxes within the violin plots show the medians as centres, interquartiles as hinges and 1.5 times the interquartile ranges as whiskers.

### 6. Data curation and analysis of the virus-infected donor dataset

This dataset was derived from a donor with prior infections of Influenza, Epstein–Barr virus (EBV), and human cytomegalovirus (HCMV), who was also diagnosed with lung cancer at the time of sample collection [4]. Peripheral blood and tumor-infiltrating T cells were isolated from the donor. Bulk sequencing of the CDR3 regions of human TCRβ chains was performed using the immunoSEQ assay. Antigen-specific T cells were obtained by co-culturing the expanded T cells with HLA-matched viral peptides. The antigen panel included Influenza A $M1_{58\text{-}66}$ (GILGFVFTL), Influenza A $PA_{46\text{-}54}$ (FMYSDFHFI), EBV $BMLF1_{280\text{-}288}$ (GLCTLVAML), and HCMV $pp65_{495\text{-}503}$ (NLVPMVATV). In total, 44 unique TCRs specific to these four viral antigens were identified. For each TCR, we computed its minimum Euclidean distance to all TCRs in the benchmark dataset. The distribution of these distances is presented in **Supplementary Fig. 4b**.

### 7. Data curation and analysis of T1D donors cohort

T cells reactive to the type 1 diabetes (T1D) antigen VAANIVLTV, derived from $ZnT8_{186\text{-}194}$, were identified using multimer (MMr) staining assays [5]. The resulting set included sixteen T cell clones—nine from five T1D patients and seven from five healthy donors. After merging clonotypes with identical TCRβ CDR3 sequences, the final dataset comprised fourteen unique clonotypes, including six from five T1D patients and eight from six healthy donors. Additionally, an MMr was designed for the *B. stercoris* WP_060386636.1-derived mimotope KAANIVLTV to evaluate potential cross-reactivity. In three out of four tested donors, double-positive $MMr^+CD8^+$ T cells reactive to both VAANIVLTV and KAANIVLTV were detected, suggesting potential cross-recognition. For each TCR and each antigen in this dataset, we computed the minimum Euclidean distance to all TCRs and antigens in the benchmark dataset. The resulting distance distributions are shown in **Supplementary Fig. 4c**.

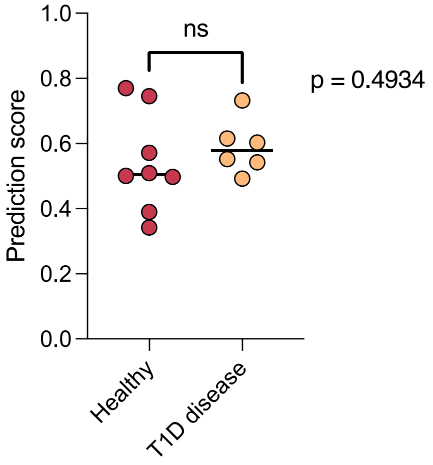


**Supplementary Fig. 6 Comparison of predicted arAg–TCR binding scores between healthy donors and T1D patients.** Each dot represents the averaged prediction score (across five cross-validated models) for a single VAANIVLTV–TCR pair. TCRs were grouped by donor disease status (healthy, n = 8; T1D, n = 6). Although the average score in the T1D group was slightly higher, the difference was not statistically significant (unpaired t-test, $p$ = 0.4934).

We then applied the five cross-validated models to predict binding probabilities for all $ZnT8_{186-194}$–TCR pairs derived from both T1D patients and healthy donors. The predicted scores from each model were averaged to obtain final binding probabilities for each TCR. As shown in **Supplementary Fig. 6**, when stratified by disease status, the average prediction scores in the T1D group were slightly higher than those in the healthy group. However, the difference was not statistically significant (unpaired t-test, $p$ = 0.4934), suggesting comparable model-inferred binding potential across groups.

### 8. Detailed explanation of antigenic immune response entropy (AIRE)

We propose antigenic immune response entropy (AIRE) as a unified metric to quantify antigen-specific immune responses by simultaneously considering clonal diversity (breadth), clonal expansion (depth), and predicted antigen–TCR binding affinity (quality). Given an antigen $a$ and a set of antigen-specific T cell clonotypes $\mathcal{T} = \{t_1, t_2, \dots, t_n\}$, the formal definition of AIRE is:

$$AIRE_a = -\sum_{i=1}^{n} (\hat{p}_{i,a} \cdot f_i) \cdot log_2(\hat{p}_{i,a} \cdot f_i)$$

where $\hat{p}_{i,a}$ is the predicted binding probability between the $i$-th T cell clonotype $t_i$ and the antigen $a$ computed by TridentTCR, and $f_i$ represents the clonal frequency of clonotype $t_i$. The product $\hat{p}_{i,a} \cdot f_i$ measures the contribution of clonotype $t_i$ to the immune response against antigen $a$. AIRE quantifies the antigen-specific immune response as a continuous Shannon entropy-like measure, reflecting the distribution and redundancy of immune activation across clonotypes. Specifically, a higher AIRE indicates a diverse and redundant clonotype distribution, reducing the risk associated with single-clonotype exhaustion or inactivation. Conversely, a lower AIRE suggests reduced immunological coverage, implying that the antigen-specific response either relies heavily on a limited set of clonotypes or has generally diminished clonal contributions.

AIRE provides stable quantification of antigen-specific immune responses across different donors, even when individual immune strategies differ substantially. For example, within the 10x Genomics dataset, both Donor 1 and Donor 2 were seropositive for the influenza A, with comparable AIRE scores ($AIRE_{Influenza}$ = 0.214 for Donor 1; $AIRE_{Influenza}$ = 0.251 for Donor 2). To further investigate these responses, we compared the clonal architectures between the two donors by identifying 38 antigen-specific clonotypes in Donor 1 and 17 clonotypes in Donor 2, and then assessing their respective contribution distributions using an unpaired t-test. As illustrated in **Supplementary Fig. 7**, the donors exhibited significantly different clonotype distributions ($p$ = 0.0003). Donor 1's repertoire was dominated by a small number of highly expanded clonotypes with high predicted binding probabilities, whereas Donor 2 showed a broader yet less expanded clonotype distribution. Both patterns indicate effective but fundamentally distinct immune-memory strategies, which AIRE captures quantitatively (see **Fig. 4** in main text).

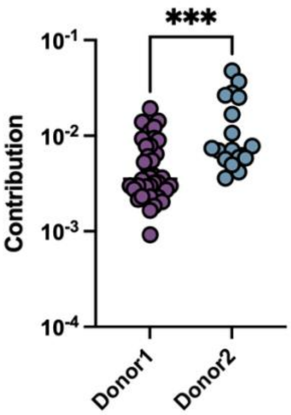


**Supplementary Fig. 7 Distinct patterns of clonotype contribution to the Influenza A $M1_{58\text{-}66}$ (GILGFVFTL) in two seropositive donors.** Each dot represents a single antigen-specific T-cell clonotype; the y-axis shows its contribution score (predicted binding probability × clonal frequency). Purple, Donor 1 (n = 38); blue, Donor 2 (n = 17). Bars denote median values. Statistical significance was evaluated with an unpaired t-test ($p$ = 0.0003).
